# The KZFP/KAP1 system controls transposable elements-embedded regulatory sequences in adult T cells

**DOI:** 10.1101/523597

**Authors:** Flavia Marzetta, Laia Simó-Riudalbas, Julien Duc, Evarist Planet, Sonia Verp, Priscilla Turelli, Didier Trono

**Affiliations:** School of Life Sciences, Ecole Polytechnique Fédérale de Lausanne (EPFL), 1015 Lausanne, Switzerland

**Keywords:** KAP1, KZFPs, transposons, endogenous retroelements, epigenetics, transcription, speciation, T lymphocytes

## Abstract

Transposable elements-embedded regulatory sequences (TEeRS) are subjected to early embryonic repression through sequence-specific recruitment of KRAB zinc finger proteins (KZFPs), their cofactor KAP1/TRIM28 and associated chromatin modifiers. This modulates the TEeRS-mediated regulation of gene expression in embryonic stem cells (ESCs) and leads to DNA methylation-induced silencing. However, KZFPs are broadly expressed in adult tissues, suggesting that they control TEeRS throughout life. Confirming this hypothesis, we reveal here that the KZFP/KAP1 system exerts a highly dynamic control of TEeRS in adult human CD4^+^ T lymphocytes. First, we observed that in these cells many TEs are still bound by KAP1, the recruitment of which is dynamically regulated upon T cell receptor stimulation. Second, we found that KAP1 depletion induces broad transcriptional alterations in T cells, with de-repression of TE-based regulatory elements leading to the illegitimate activation of nearby genes. Finally, we show that the tissue-restricted expression of KZFPs correlates with KAP1-mediated lineage-specific chromatin signatures and transcriptional repression. These data support a model where TE-targeting KZFPs and KAP1 are important regulators of gene expression in adult human cells.

## INTRODUCTION

More than half of the human genome is readily recognizable as derived from transposable elements (TEs), mostly endogenous retroelements (EREs) such as long terminal repeat (LTR) retrotransposons, also known as endogenous retroviruses (ERVs), long and short interspersed elements (LINEs and SINEs, respectively), or SINE-VNTR-Alu (SVAs) (Lander et al. 2001). Transposons were once described as purely parasitic or junk DNA, maintained within genomes solely due to their replicative capacity (Doolittle and Sapienza 1980; Orgel and Crick 1980). However, cumulated evidence indicates that they significantly contribute to genome evolution by introducing genetic variation through multiple mechanisms (Beck et al. 2010; Huang et al. 2010; Iskow et al. 2010). For instance, TEs can provide alternative promoters (Faulkner et al. 2009), splice sites (Lin et al. 2008) and polyadenylation signals (Lee et al. 2008) and can disseminate transcription factor binding sites hence participate in the regulation of gene expression (Jacques et al. 2013; Fort et al. 2014; Sundaram et al. 2014; Pavlicev et al. 2015). While the evolutionary benefits of these effects emerge through natural selection, TEs can also be acutely detrimental, for instance when new integrants inactivate open reading frames or severely disrupt gene expression, and retrotransposons from all groups still active in humans have been implicated in the genesis of disease (Kaer and Speek 2013). Nevertheless, only a very small fraction of human TEs, about 1 in 10,000, is still competent form transposition, all the others being incapacitated in this function by mutations or deletions (Friedli and Trono 2015).

As genomic threats, TEs, whether or not transposition-competent, are subjected to tight control, which both represses their transcription and, if relevant, prevents their genomic spread (Rowe and Trono 2011; Schlesinger and Goff 2015). While DNA methylation at CpG dinucleotides is thought to represent a stable and long-lasting mechanism of ERE silencing (Reik 2007), histone-based repression appears essential during the phases of genome reprogramming that take place in early embryos (Morgan et al. 2005; Karimi et al. 2011). KRAB-containing zinc finger proteins (KZFPs), which are encoded in the hundreds by the mouse and human genomes (Emerson and Thomas 2009), are key mediators of this process. Endowed with sequence-specific DNA binding ability through arrays of zinc fingers, KZFPs recruit to TEs their cofactor KAP1 (also known as TRIM28 or TIF1β) (Friedman et al. 1996), which coordinates the assembly of a macromolecular complex containing chromatin-modifying enzymes such as the histone 3 lysine 9-specific methyltransferase SETDB1 (Schultz et al. 2002) and the histone deacetylase contained in the NuRD complex (Schultz et al. 2001) as well as heterochromatin protein 1 (HP-1) (Lechner et al. 2000) and, at least during early embryonic development, DNA methyltransferases (Wiznerowicz et al. 2007; Quenneville et al. 2012). These effectors cooperatively induce the formation of heterochromatin thus repress transcription through epigenetic changes (Sripathy et al. 2006).

Depleting KAP1 in mouse or human embryonic stem cells (ESCs) leads to the transcriptional activation of a wide spectrum of TE-derived loci through the loss of SETDB1-dependent H3K9 trimethylation and other repressive marks (Matsui et al. 2010; Rowe et al. 2010; Castro-Diaz et al. 2014; Jacobs et al. 2014; Turelli et al. 2014). It also perturbs the transcriptional equilibrium of these cells notably by unleashing the activity of TE-embedded regulatory sequences (TEeRS), such as ERE-based promoters and enhancers impacting on the expression of nearby genes (Rowe et al. 2013b; Turelli et al. 2014). In contrast, deleting KAP1 in mouse embryonic fibroblasts was found to induce comparatively modest transcriptional deregulations (Rowe et al. 2010; Rowe et al. 2013b). This was interpreted as reflecting the prior DNA methylation of TEeRS, as this self-perpetuating silencing mark alleviates the need for persistent expression of sequence-specific repressors (Wiznerowicz et al. 2007; Quenneville et al. 2012; Rowe et al. 2013a). However, conditional *KAP1* knockout in a variety of mouse organs systematically led to abnormal phenotypes, linking the master regulator with processes as diverse as erythropoiesis (Barde et al. 2013), B and T lymphocyte activation and development (Chikuma et al. 2012; Santoni de Sio et al. 2012a; Santoni de Sio et al. 2012b), hepatic metabolism (Bojkowska et al. 2012), skeletal muscle differentiation (Singh et al. 2015) and management of behavioral stress (Jakobsson et al. 2008). While KAP1 is known to accomplish some effects in a KZFP and TE-independent manner (Iyengar et al. 2011; Singh et al. 2015; McNamara et al. 2016), mounting evidence indicates that TEeRS and their cognate controllers regulate the physiology of higher vertebrates beyond the early embryonic period. First, the KZFP/KAP1 system controls a subset of LTR-retrotransposons and neighboring genes in mouse neuronal progenitors (Fasching et al. 2015) and multiple adult cell types (Ecco et al. 2016; Tie et al. 2018). Second, SETDB1-dependent H3K9me3 is required to repress endogenous retroviruses in pro-B lymphocytes of adult mice (Collins et al. 2015). Third, most KZFPs have TEs as prominent genomic targets (Najafabadi et al. 2015; Imbeault et al. 2017), yet many are expressed in adult tissues both in human and mouse (Lizio et al. 2015; Imbeault et al. 2017). Finally, a significant fraction of TEs bound by KAP1 in human ESCs still bear the corepressor in peripheral blood T lymphocytes (Turelli et al. 2014). The present study was undertaken to explore beyond the early embryonic period the influence of KZFPs and their TE targets on transcriptional regulation networks. Using CD4^+^ T lymphocytes as a model system, we demonstrate that the KZFP/KAP1-mediated control of TEeRS sharpens the lineage-specificity and activation-responsiveness of gene expression in an adult human tissue.

## RESULTS

### KZFP/KAP1-mediated control of EREs is maintained in CD4^+^ T lymphocytes

We used chromatin immunoprecipitation followed by deep sequencing (ChIP-seq) to investigate KAP1 recruitment to repetitive sequences in human peripheral blood CD4^+^ T lymphocytes. We found that a significant fraction of KAP1 ChIP-seq peaks detected on EREs in ESCs was maintained in CD4^+^ T cells (Fig. 1A). KAP1 recruitment at these genetic units did not simply reflect their overall abundance in the genome, with for instance enrichment at specific HERV (human ERV) and SVA integrants, and was very significantly modified in response to T cell receptor (TCR) stimulation (Fig. 1B; Supplemental Fig. S1).

**Figure 1.**
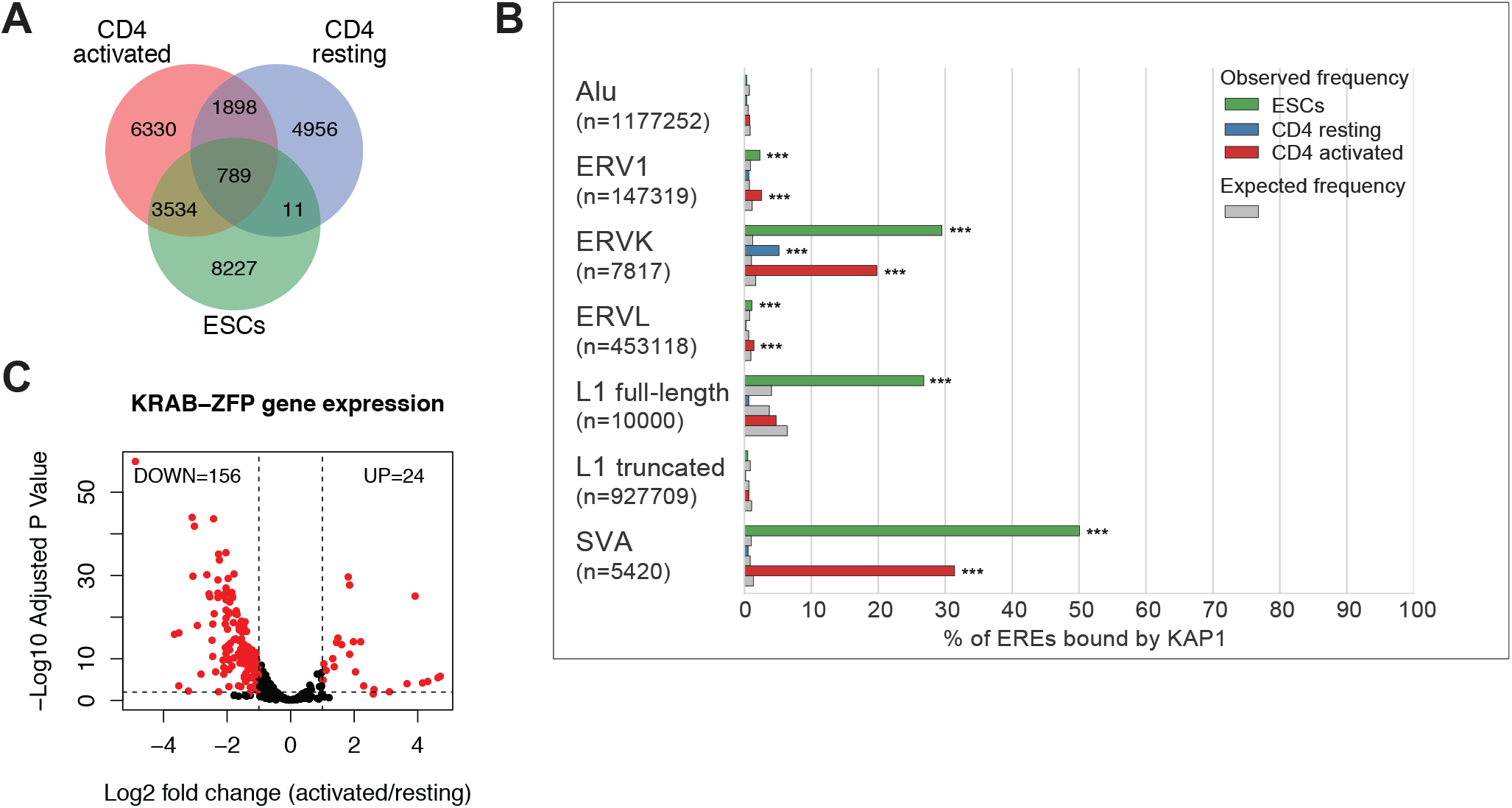
KAP1 binding to EREs is maintained in adult CD4^+^ T cells. (A) Venn diagrams depicting overlap between KAP1 ChIP-seq peaks on EREs commonly detected in two replicate ChIP-seq experiments performed on chromatin from human H1 ESCs, resting or activated CD4^+^ T lymphocytes. (B) Bar plot showing the percentage of ERE integrants on which we detected KAP1 ChIP-seq peaks in H1 ESCs, resting and activated CD4^+^ T cells. For full-length L1 integrants, a length cutoff of 5kb was used. For each family, both the values observed (color bars) or expected (grey bars) are shown. Expected frequency and statistical significance was calculated using Bedtools Fisher’s test. Similar results were obtained from independent ChIP-seq experiments (Supplemental Fig. 1A). (C) Volcano plot displaying KZFP genes expressed in activated and resting CD4^+^ T lymphocytes. Colored data points depict genes with a significant expression change (> 2-fold where P_adj_ < 0.05).

KAP1 is recruited to TEs via KZFPs, the majority of which bind specific subsets of these elements (Najafabadi et al. 2015; Imbeault et al. 2017). Correspondingly, we found many of these proteins to be expressed in CD4^+^ T lymphocytes, with different levels in resting and activated cells as determined by deep RNA sequencing (RNA-seq) (Fig. 1C).

### TCR signaling influences the dynamics of KAP1 genomic recruitment in CD4^+^ T cells

Overall, about 90, 50 and 70% of KAP1 peaks detected in ESCs, resting and activated CD4^+^ T cells, respectively, were found to overlap with EREs (Turelli et al. 2014). To gain functional insight, we compared the genomic context of KAP1 recruitment in resting and activated T lymphocytes. KAP1 binding sites restricted to the quiescent CD4^+^ T cell state were evenly distributed in and out of annotated EREs and at least half of them resided within less than 10kb of the 5’ end of a gene (Fig. 2AB). In contrast, genomic sites where KAP1 was *de novo* recruited in activated CD4^+^ T cells were generally located at greater distances from transcriptional start sites (TSS) and mostly coincided with EREs (Fig. 2AB). *KZFP* gene loci, previously observed to be bound by ZNF274 and KAP1 in several cell types (Frietze et al. 2010; Iyengar et al. 2011), recapitulated some of these shifting distribution patterns. In resting cells, they preferentially harbored KAP1 at their promoter, whereas upon activation the regulator was displaced to their 3’, ZNF array-coding exon (Supplemental Fig. S2).

**Figure 2.**
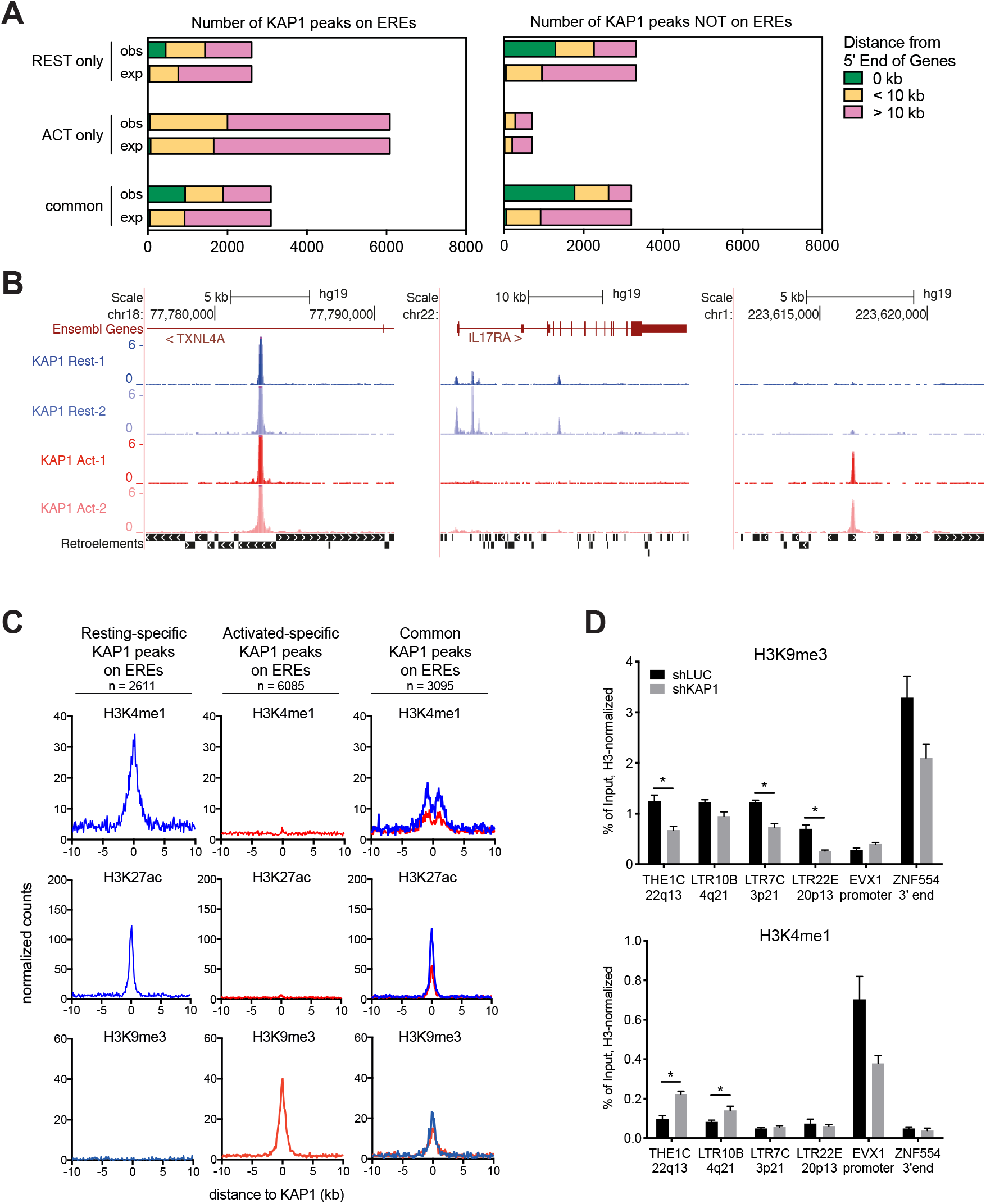
KAP1 genomic recruitment is widely redistributed following T cell activation. (A) Number of KAP1 ChIP-seq peaks located at the 5’ end of genes, within 10kb or at further distance. Left, KAP1 peaks overlapping with EREs. Right, KAP1 peaks not overlapping with EREs. KAP1 peaks were grouped according to whether they were detected exclusively in resting (REST only) or activated (ACT only) CD4^+^ T cell ChIP-seq experiments or in all data sets (common). As a control, we plotted the distribution of KAP1 peaks that we obtained by averaging a 1000-time permutation of random assignation of KAP1 peak coordinates. (B) UCSC Genome Browser snapshots of KAP1 ChIP-seq profiles providing example of conserved or differential binding between the resting and activated T cell conditions. Results of replicate experiments are shown. (C) Positional relationship between KAP1 peaks on EREs and the indicated histone marks, at a 100-bp resolution and over a 10kb window centered on KAP1 ChIP-seq peaks. Left, KAP1 peaks detected exclusively in resting cells. Middle, KAP1 peaks detected exclusively in activated cells. Right, KAP1 peaks detected in both datasets. The profiles were normalized for the total number of ChIP-seq peaks in each sample. (D) H3K9me3 and H3K4me1 ChIP-qPCR analysis in control and *KAP1* knockdown cells at selected KAP1-bound ERE integrants. Values are normalized to their respective total inputs and to the total H3 levels. *EVX1* and *ZNF554* 3’ were chosen, respectively, as control regions for H3K4me1 and H3K9me3. Error bars represent SEM of biological replicates (n=3), *p < 0.05, **p < 0.01, ***p < 0.001, paired t test.

To examine chromatin features associated with KAP1 genomic recruitment, we used a combination of our and previously published ChIP-seq data from resting or activated CD4^+^ T cells (Bernstein et al. 2010; Hawkins et al. 2013). In quiescent lymphocytes, KAP1 recruitment strongly correlated with the presence of H3K4me1 and H3K27ac, a combination typically found at active enhancers and promoters (Creyghton et al. 2010; Calo and Wysocka 2013) (Fig. 2C and Supplemental Fig. S3, left panels). Upon T cell activation, KAP1 peaks were frequently detected in association with the repressive mark H3K9me3, especially when they coincided with EREs (Fig. 2C and Supplemental Fig. S3, middle panels). Interestingly, at some KAP1-recognized ERE loci (243 and 144 in resting and activated cells, respectively), not only was the binding of the corepressor dynamic between the two cell states, but it also coincided with the repressive H3K9me3 mark in one condition and the active H3K27ac mark in the other (Supplemental Fig. S4A). This suggests that KAP1-controlled TEeRS could act as T cell activation-dependent regulatory elements, a model also supported by their frequent proximity to gene promoters (Supplemental Fig. S4B).

### KAP1 is required to maintain repressive chromatin during T cell activation

We next explored the consequences of depleting KAP1 on the chromatin status of activated T lymphocytes. For this, 24h after CD3/CD28-stimulation, CD4^+^ cells were transduced with lentiviral vectors (LV)-expressing small hairpin RNAs (shRNAs) targeting either *KAP1* or *Luciferase* and encoding for green fluorescent protein (GFP). Five days later, KAP1 depletion exceeded 90% in GFP-positive cells expressing the corresponding shRNA, as assessed by immunoblotting (Supplemental Fig. S5). Using chromatin immunoprecipitation and quantitative PCR (ChIP-qPCR), we examined the consequences of KAP1 depletion at ERE integrants previously defined as enriched for this regulator in activated cells (Supplemental Fig. S6). At these loci, *KAP1* knockdown induced a drop in H3K9me3, often matched by a gain in H3K4me1 (Fig. 2D). Taken together, these results reveal a functional link between KAP1 and the maintenance of epigenetic repression during T cell activation, notably at EREs.

### KAP1 depletion perturbs the CD4^+^ T cell transcriptome

In embryonic stem cells and some differentiated cell lines, KAP1-mediated epigenetic repression of TEeRS influences the expression of nearby genes (Rowe et al. 2013b; Turelli et al. 2014; Ecco et al. 2016). By analogy, we hypothesized that TEeRS and their epigenetic controllers might condition the transcriptional dynamics of CD4^+^ T cells. To probe this issue, we subjected KAP1-depleted TCR-stimulated T lymphocytes to RNA-seq. To minimize false positives due to off-target effects, we used two different *KAP1*-targeting shRNAs and considered only transcriptional changes induced by both of them (Fig. 3A). Based on a twofold cut-off and a P_adj_ < 0.05, we identified more than 2,000 EREs, the expression of which differed in *KAP1* knockdown compared to control cells (Fig. 3B, left panel). Most were upregulated, and even though KAP1 was detected at baseline at only a fraction of these EREs, they were proportionally more KAP1-bound than their transcriptionally unchanged counterparts (Fig. 3C, left panel). *KAP1* knockdown also induced extensive changes in the gene-derived transcriptome of activated T cells (Fig. 3B, right panel). However, the spectrum of these transcriptional changes was broader, with up-regulated genes accounting for only 40% of a total of about 1,400 perturbed units. Furthermore, genes induced upon KAP1 depletion did not bind the repressor more often than genes that remained stable in this setting (Fig. 3C, right panel). Nevertheless, we observed a positive correlation between expression of EREs and nearby genes upon KAP1 depletion (Fig. 3D). Importantly, the correlation between EREs and genes was independent of their respective orientations, ruling out the possibility that ERE expression was merely due to read-through transcripts (Supplemental Fig. S7). Overall, these observations indicate that KAP1 partakes in the epigenetic control of EREs in CD4^+^ T cells and that disruption of this process leads to global transcriptional alterations.

**Figure 3.**
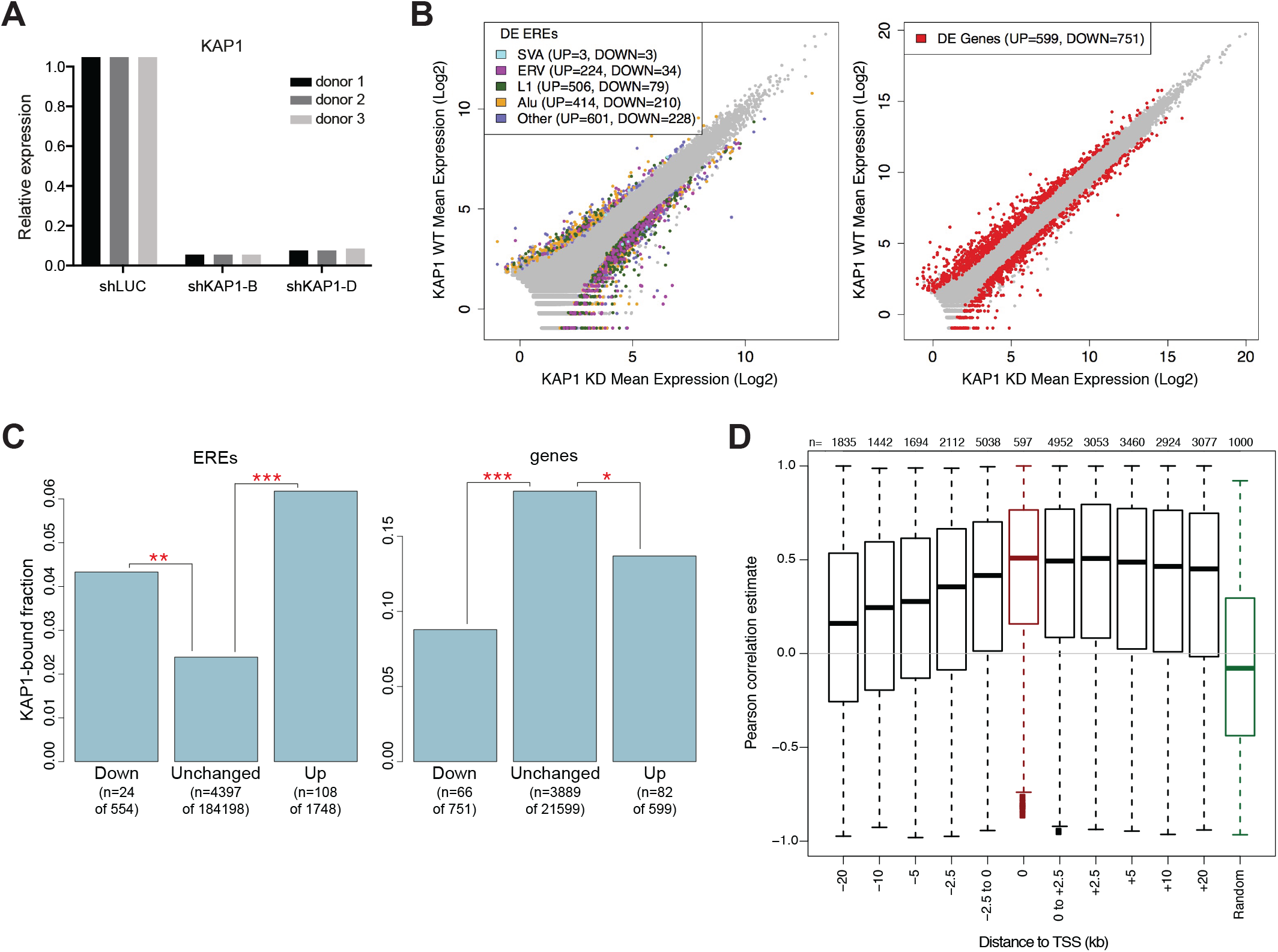
KAP1 depletion broadly perturbs CD4^+^ T cells transcriptional dynamics. (A) RT-qPCR analysis of KAP1 downregulation in RNA samples from activated CD4^+^ T cell subjected to high-throughput sequencing. (B) Comparison of ERE (left panel) or gene (right panel) expression data from *KAP1* knockdown (shKAP1) and control (shLUC) activated CD4^+^ T cell. Results are presented as average normalized counts for each condition. Each ERE integrant or gene is represented by a single data point. Colored data points indicate a significant (P_adj_ < 0.05) and greater than two-fold change in expression. (C) Proportion of EREs (left panel) or genes (right panel) that are KAP1-bound within each of the three groups (i.e. downregulated, unchanged and upregulated in *KAP1* knockdown compared to control cells). The proportion of bound elements was tested using a Fisher’s exact test. (D) Pearson correlation coefficients between expressed EREs and closest genes. EREs were grouped into several bins of distance to genes, further differentiating between those located upstream or downstream the genes. EREs overlapping with gene introns are highlighted in red.

We similarly attempted to assess the impact of KAP1 depletion in quiescent T cells (Supplemental Fig. S8). The levels of knockdown obtained in this setting, even after transduction of cells transiently stimulated with IL-7 to increase their permissiveness to LV-mediated transduction (Ducrey-Rundquist et al. 2002), were lower (~80% compared with >95% in activated cells). Nevertheless, changes in both EREs and genes expression were very negligible compared to their activated counterparts. It is possible that in quiescent T cells KAP1 plays a less pronounced role in epigenetic silencing, as reflected by its restricted association with the repressive H3K9me3 mark (Fig. 2C). Additionally, the consequences of KAP1 depletion on the CD4^+^ T cells transcriptional dynamics might emerge only after few rounds of cell division, when the repressive mark get effectively diluted.

### The KZFP/KAP1-mediated control of TEeRS preserves the activation-responsiveness and lineage-specificity of CD4^+^ T cell gene expression

To attempt identifying the events physiologically regulated by EREs and their controllers, we first monitored the *ex vivo* expansion of KAP1-depleted, GFP-positive cells in response to CD3/CD28 stimulation. *KAP1* knockdown cells significantly decreased in number over time compared with control cells (Supplemental Fig. S9A). In agreement with a loss of cell viability, we detected an increase in surface exposure of the apoptotic marker Annexin V (Supplemental Fig. S9B). Furthermore, the cell cycle profile of *KAP1* knockdown cells revealed an incomplete progression to G2M (Supplemental Fig. S9C), indicating a defective proliferation in response to activating stimuli.

To explore the molecular tenets of these phenotypic features, we compared the transcriptional alterations induced by KAP1 depletion in activated CD4^+^ T cells with changes in gene expression that occur upon activation of resting cells (Supplemental Fig. S9D). Out of 4,766 genes normally induced by TCR stimulation, the expression of 328 genes was instead decreased if KAP1 was depleted. Conversely, KAP1 loss caused the upregulation of 243 of the 5,356 genes, the expression of which was normally attenuated following T cell activation. This indicates that the loss of KAP 1 impairs, directly or indirectly, transcriptional patterns induced by TCR stimulation, impacting on T cell homeostasis.

We next examined whether KAP1 control over TEeRS could contribute to establish tissue-specific patterns of gene expression in CD4^+^ T cells. To probe this possibility, we selected a few genes upregulated in *KAP1* knockdown T lymphocytes, which had their TSS situated within 10kb of a KAP1-bound ERE and were differentially expressed across cell types. Indeed, we previously determined that KAP1-nucleated heterochromatin can readily spread over this distance (Groner et al. 2010); furthermore, promoter-enhancer interactions are more likely and easier to identify over short distances. We focused on one of these genes, *TLR1*(toll-like receptor 1), which encodes for an innate immune receptor recognizing pathogen-associated molecular patterns and is expressed at higher levels in myeloid than in lymphoid cells (Ochoa et al. 2003). In CD4^+^ T cells, we found KAP1 bound to an ERV-derived sequence (*MER50, LTR18B*) located in an intron approximately 2kb downstream of the *TLR1* promoter (Fig. 4A). Some H3K27Ac and, to a lesser extent, H3K4me1 were detected at the promoter of this gene, but its transcribed region was devoid of the active chromatin markers and instead covered with H3K9me3, especially at the KAP1-bound ERV (Fig. 4A). Differences in *TLR1* expression between lymphoid and myeloid tissues correlated with distinctive epigenetic features. In neutrophils, where this gene is highly expressed (Forrest et al. 2014), it was devoid of H3K9me3, including over the *MER50* and *LTR18B* sequences, and instead markedly enriched in the active marks H3K4me1, H3K4me3 and H3K27Ac (Fig. 4A). In KAP1-depleted T lymphocytes, H3K9me3 dropped at the intronic ERV whereas H3K27Ac accumulated at the nearby promoter (Fig. 4B), which correlated with increased transcription (Fig. 4AC). Finally, we examined the impact of KAP1 depletion on *TLR1* expression in HL-60 cells, a promyelocytic cell line that can be induced to differentiate with DMSO (Jacob et al. 2002). *TLR1* RNA levels were higher in HL-60 than in CD4^+^ T cells, as predicted, yet this gene was strongly induced by DMSO treatment (Fig. 4D). Most interestingly, *TLR1* was activated by KAP1 depletion in untreated HL-60 cells, but the knockdown had no further effect in their DMSO-differentiated counterparts (Fig. 4E). A similar, KAP1-dependent switch from repressive to active histone marks was observed in proximity of other derepressed genes that displayed sustained expression and active chromatin configuration in hematopoietic lineages different from CD4^+^ T cells (Supplemental Fig. S10).

**Figure 4.**
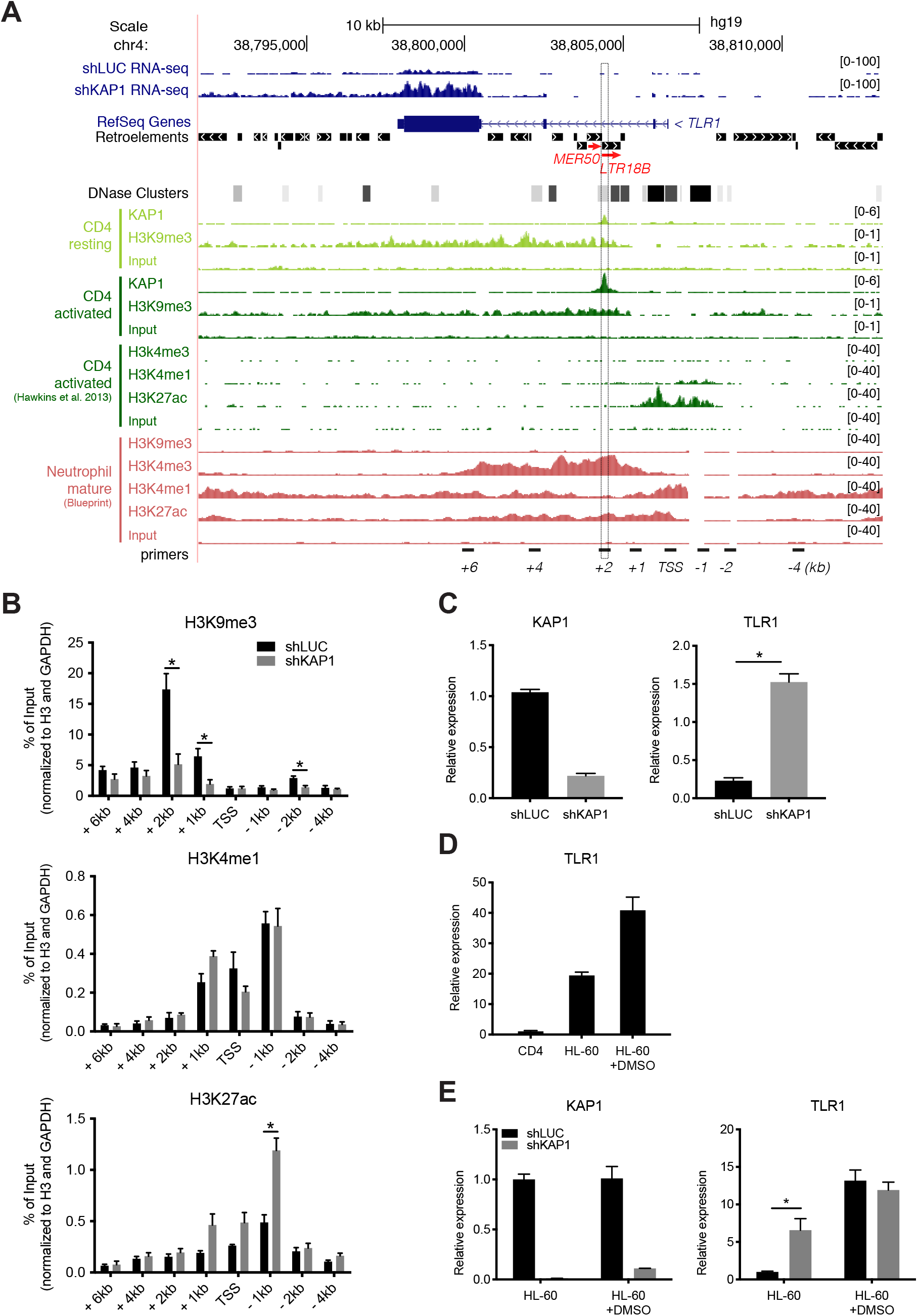
KAP1–mediated control of a ERE-based regulatory element modulates the lineage-specific expression of *TLR1*. (A) UCSC Genome Browser view of *TLR1* locus. Displayed tracks include RNA-seq for control (shLUC) and *KAP1* knockdown (shKAP1) activated CD4^+^ T cells; RefSeq genes and retroelement annotation (modified from UCSC Genome Browser as described in the Methods); DNase cluster (ENCODE); KAP1 and H3K9me3 ChIP-seq tracks for resting and activated CD4^+^ T cells; public H3K9me3, H3K4me3, H3K4me1, and H3K27ac ChIP-seq tracks for Th1 activated CD4^+^ T cells and/or mature neutrophils; primers used in (B) located at varying distances relative to the TSS. The dashed box highlights the KAP1-bound region. The repetitive elements on which KAP1 binding is centered are shown in red. (B) ChIP-qPCR analysis of H3K9me3, H3K4me1 and H3K27ac around KAP1 binding site at the *TLR1* locus in control and *KAP1* knockdown cells. Values are normalized to their respective total inputs, to the total H3 protein levels and to *GAPDH*. (C) RT-qPCR analysis of *KAP1* and *TLR1* mRNA expression in activated CD4^+^ T lymphocytes transduced with shLUC or shKAP1-expressing LVs. (D) RT-qPCR analysis of *TLR1* mRNA expression in activated CD4^+^ T lymphocytes and in the human promyelocytic leukemia cell line HL-60. HL-60 cells were differentiated into neutrophil-like cells in culture medium in the presence of 1.3% DMSO for five days. (E) RT-qPCR analysis of *KAP1* and *TLR1* mRNA expression in untreated or DMSO-treated HL60 cells, transduced with shLUC or shKAP1-expressing LVs. Error bars represent SEM of biological replicates (n=3), *p < 0.05, **p < 0.01, ***p < 0.001. Paired t test was used in (B) and (C) and unpaired t test was used in (E).

### CD4-specific expression of KZFPs contributes to tissue-specific regulation of gene expression

Even though we could not identify the KZFP responsible for recruiting KAP1 at the *TLR1/MER50-LTR18B* locus, we reasoned that a CD4-restricted expression of KZFPs could contribute to lineage-specific chromatin signature and transcriptome. To test this hypothesis, we selected KZFPs peaks identified in 293Ts by chromatin immunoprecipitation with exonuclease digestion (ChIP-exo) (Imbeault et al. 2017) that were located within EREs and computed the overlap between these repetitive elements and H3K9me3, H3K4me1 and H3K27ac histone mark profiles of 50 cell types from the NIH Roadmap database (Roadmap Epigenomics et al. 2015). Results indicated that the associations between KZFP-bound EREs and histone marks varied depending on the recruited protein, as well as on the cell type (Fig. 5A and Supplemental Fig. S11). Furthermore, consistent with our previous observations (Imbeault et al. 2017), not all KZFP binding sites on TEs displayed statistically significant association with repressive chromatin, supporting the hypothesis of a plurality of functions for this family of transcriptional modulators.

**Figure 5.**
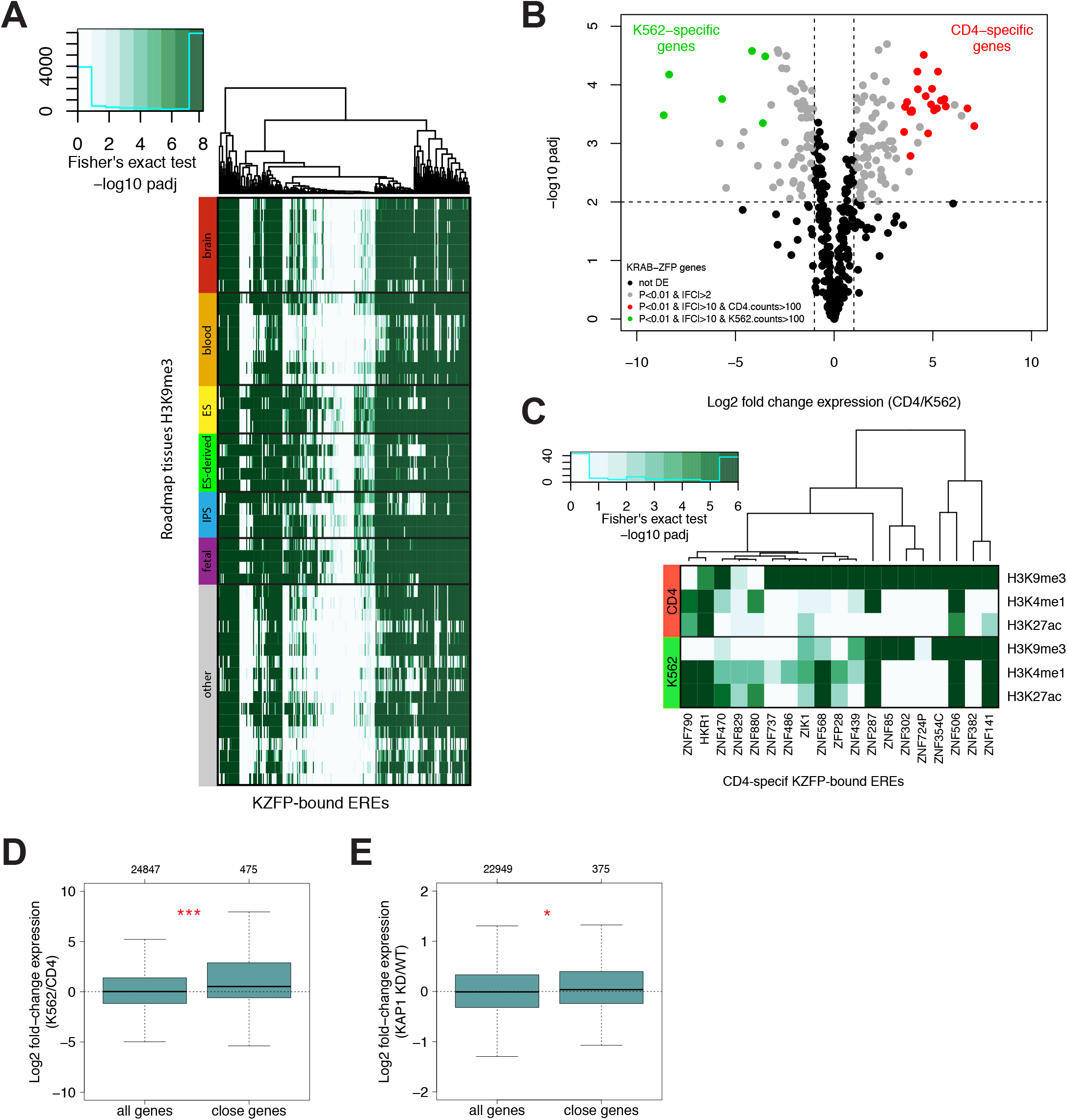
CD4-specific KZFPs contribute to tissue-specific regulation of gene expression. (A) Heatmap showing the Fisher’s Exact test adjusted P-values for the overlap between KZFP-bound EREs (X-axis, where each column represents a different KZFP ChIP-exo sample) and H3K9me3 from the NIH roadmap dataset (Y-axis). The score is -log10 of Benjamini-Hochberg-corrected P-values. (B) Volcano plot displaying KZFP genes expressed in activated CD4^+^ T lymphocytes and K562 cells based on RNA-seq data. Red and green data points depict genes with a significant higher expression in CD4^+^ or K562 cells, respectively (> 100 normalized counts and > 10-fold changes where P_adj_ < 0.01). (C) Similar to (A), but KZFPs on the X-axis are selected for their CD4-specific expression and histone modifications on the Y-axis represents activated CD4^+^ T lymphocytes and K562 dataset. (D) Boxplots showing fold-change expression of all genes or genes whose TSS is located within 10kb from “CD4-specific” KZFP-bound EREs that were marked by H3K9me3 in CD4^+^ cells and by H3K4me1 and H3K27ac in K562 cells, in K562 compared to activated CD4^+^ T cells. (E) Similar to (D), but fold-change expression of genes is relative to KAP1-depleted and control activated CD4^+^ T cells. *p < 0.05, **p < 0.01, ***p < 0.001, t test.

We then decided to focus on a subset of KZFPs that, based on their expression profiles, could act as tissue-specific transcriptional regulators in CD4^+^ T cells. We compared the *KZFP* gene transcriptome of activated CD4^+^ T lymphocytes and K562 leukemia cells, for which both RNA-seq and histone marks profiles are publicly available (Pradeepa et al. 2016; Liu et al. 2017), and we selected genes that showed strong and specific expression in CD4^+^ cells (Fig. 5B). The majority of EREs bound by CD4-specific KZFPs were significantly associated with repressive chromatin in CD4^+^ T lymphocytes but less frequently in K562 cells, where their chromatin profile was more diversified (Fig. 5C).

Finally, to determine whether CD4-specific KZFP-bound TEeRS could serve as tissue-specific regulators for gene expression, we selected those elements that were marked by H3K9me3 in CD4^+^ T cells and by H3K4me1 and H3K27ac in K562 cells. We then analyzed the expression of genes, the TSS of which was located within 10kb upstream or downstream these loci. This revealed that genomic proximity to these repetitive elements matched with higher gene expression in K562 compared to activated CD4^+^ T cells (Fig. 5D). Additionally, these genes were significantly up-regulated in *KAP1* knockdown CD4^+^ T cells, as expected if KZFPs act through KAP1 to exert a tissue-specific control of TEeRS and their target genes (Fig. 5E). Altogether, these data support a model whereby the control of TEeRS by the KZFP/KAP1 system contributes to the lineage- and activation-specificity of gene expression in adult human cells.

## DISCUSSION

TEeRS are subjected to dynamic control in germ cells and pre-implantation embryos, where they display highly stage-specific patterns of expression and impact on the regulation of host genes (Rowe and Trono 2011). During this early developmental period, they are progressively repressed by tightly orchestrated epigenetic mechanisms mediated by KZFPs, their cofactor KAP1 and associated chromatin modifiers (Matsui et al. 2010; Rowe et al. 2010; Castro-Diaz et al. 2014; Jacobs et al. 2014; Turelli et al. 2014). This leads to their DNA methylation, a modification thought to induce their permanent silencing, with re-activation only in exceptional circumstances such as during the late phase of neuronal stem cell differentiation or in some cancers (Coufal et al. 2009; Iskow et al. 2010). However, many enhancers active in adult tissues reside within TEs, which explains why the genomic locations of many transcription factor binding sites are species-specific (Chuong et al. 2016). Our work suggests that the KZFP/KAP1 system modulates the activity of TEeRS to shape cell-restricted gene expression in human adult T lymphocytes. These results extend our earlier description of KZFPs controlling ERE-mediated gene regulation in mouse adult tissues both *in vitro* and *in vivo* (Ecco et al. 2016).

We were first puzzled upon finding that numerous EREs bound by KAP1 in human ESCs still bore the corepressor in peripheral blood CD4^+^ T lymphocytes, and that these cells expressed many *KZFPs* known to target EREs. By comparing KAP1 genomic recruitment in resting and activated lymphocytes, we went on to uncover a dynamic picture, which impacted on the expression of both EREs and cellular genes. In resting cells, KAP1 was preferentially recruited at transcription start sites and regions adorned with active chromatin marks. Upon T cell activation, its enrichment at these loci was dampened, and it was displaced towards H3K9me3-rich regions, most of which were EREs, coincident with a strengthening of the repressive mark at these loci. This confirms that KAP1 can associate with different functional complexes, with contrasting transcriptional outcomes. The marked presence of KAP1 at TSS of active genes, notably in resting T cells, is consistent with the demonstration that it mediates the recruitment of the 7SK snRNP complex and P-TEFb to these loci, to facilitate transcription elongation by RNA polymerase II (McNamara et al. 2016). It could be that this effect is more important in the transcriptionally less active environment of resting cells, thus explaining why KAP1 vanished from many promoters following TCR stimulation. Conversely, in the latter setting it is the corepressor function of KAP1 that dominated, leading to the de novo deposition or strengthening of the repressive mark H3K9me3 at many loci, notably transposable elements. As a corollary, depleting KAP1 in activated T lymphocytes induced a rapid loss of H3K9me3 at EREs, thousands of which became transcribed. Furthermore, changes in gene expression were noted, which statistically correlated with proximity to transcriptionally perturbed EREs. In some cases, we could document that unleashing KAP1-dependent repression on EREs impacted on the expression of nearby genes in a tissue-specific fashion. For instance, a KAP1-bound LTR retrotransposon located two kilobases downstream of the *TLR1* TSS served as a seed for H3K9me3 deposition spreading over the promoter, thus avoiding that this gene, the expression of which is myeloid-restricted, be induced upon TCR stimulation. Extending this observation, we found that a large fraction of genes upregulated upon KAP1 depletion in activated T lymphocytes were normally expressed in other tissues, be it other blood cells or unrelated organs such as brain, liver or pancreas (Supplemental Fig. S12). This suggests that TEeRS and their KZFP ligands partake in maintaining cell- and tissue-specific patterns of gene expression.

By keeping a dynamic control over TEeRS beyond the early embryonic period, higher vertebrates are endowed with a reservoir of regulatory elements that can be differentially utilized, for instance through the tissue-specific expression of KZFPs or post-translational modifications altering the functional properties of either these proteins or their cofactor KAP1 (Singh et al. 2015). Our preliminary profiling of the human KZFPs interactome also indicates that these proteins can associate with a variety of complexes involved in either transcriptional or post-transcriptional regulation. Furthermore, it is likely that the range of transcriptional influences exerted by TEeRS is far more diversified than the relatively short distance cis-acing enhancer, promoter or repressor effects documented here and in other studies, whether in physiological or abnormal situations such as following KZFP, KAP1 or SETDB1 depletion (Faulkner et al. 2009; Rebollo et al. 2012a; Rebollo et al. 2012b; Rowe et al. 2013b; Turelli et al. 2014; Collins et al. 2015; Ecco et al. 2016). Longer-range influences, for instance on genome architecture or via ERE-produced regulatory RNAs (Kapusta et al. 2013; Ha et al. 2014), are indeed difficult to analyze when they involve repetitive sequences.

By exerting their influence on gene expression, TEeRS and their KZFP controllers contribute to the establishment and rewiring of species-specific regulatory networks. Indeed, TEs contribute massively to the diversification of genomes during evolution (Khan et al. 2006; Roy-Engel et al. 2008). It follows that their sequence and genomic location are poorly conserved between humans or close relatives and other species. Accordingly, a substantial fraction of KZFPs has evolved as primate-specific genes (Nowick et al. 2010; Liu et al. 2014). Therefore, the regulatory effects of TEeRS and their cognate controllers on human gene expression are evolutionary linked, and models from distant species, while certainly valid for delineating canonical aspects of gene regulation, do not allow the analysis of fine-tuned aspects of human biology.

## METHODS

### Lentiviral Vectors

For *KAP1* knockdown experiments, shRNAs against *KAP1* (shKAP1-B and shKAP1-D) were cloned into the pLKO.1.puro vector obtained from Addgene (http://www.addgene.org) using AgeI and EcoRI sites. The vectors were further modified to express a GFP reporter instead of puromycin resistance gene. The shRNA targeting sequences were obtained through the RNAi Consortium (http://www.broadinstitute.org/rnai/public). An shRNA targeting *Luciferase* (shLUC) was cloned as a control (Amendola et al. 2009). All the sequences are listed in the Supplemental Material (Supplemental Table 1). Vectors were produced by transient transfection of 293T cells detailed at http://tronolab.epfl.ch.

### Isolation, stimulation, and culture of CD4^+^ T lymphocytes

Peripheral blood mononuclear cells were isolated from buffy coat from adult healthy donors by the standard method of density gradient centrifugation using Ficoll-Paque PLUS. CD4^+^ T cells were enriched by negative selection using the EasySep Human CD4^+^ T Cell Enrichment Kit (StemCell Technologies). The purity of the CD4^+^ T cells was evaluated by flow cytometry and was higher than 95%. Cells were cultured in Xvivo 15 medium (Lonza), supplemented with 10% fetal calf serum, penicillin and streptomycin at 5 mM (Gibco), and 2 mM glutamine (Gibco) at 37°C with 5% CO2. Activation was performed by addition of Dynabeads Human T-Activator CD3/CD28 (25 ul/0.5 million cells; Life Technologies), along with IL-2 (30 U/ml; Sigma). When indicated, cells were alternatively treated with IL-7 (25 ng/ml; Lubioscience). Cells were cultured at a density of 1.0 x 10^6^ to 2.0 x 10^6^ cells/ml for the indicated time. *KAP1* knockdown was induced by transduction with shRNA vectors. Transductions of CD3/CD28-stimulated cells were done at a multiplicity of infection (MOI; determined in 293T cells) of 20, approximately 24h after addition of the beads. Transductions of IL-7-treated cells were done at MOI 40, after four days of cytokine exposure.

### Flow cytometry

To assess the purity of the population, freshly isolated CD4^+^ T cells were stained with monoclonal antibodies against CD3-PE and CD4-FITC (Biolegend) using standard procedures. The number of CD4^+^ T cells was enumerated during *ex vivo* expansion using CountBright beads (LifeTechnologies). Apoptosis was determined by Annexin V staining (Biovision). Cell cycle profile was analysed by propidium iodide staining (Invitrogen) as described by Riccardi and Nicoletti (Riccardi and Nicoletti 2006). To analyse DNA and RNA content, samples were stained with HOECHST 33342 (Sigma) and pyronin Y (Polysciences) as previously described by Ducrey-Rundquist et al. (Ducrey-Rundquist et al. 2002). Flow cytometry was performed on CyAn (Dako) or LSR II (BD biosciences) flow cytometers and analysed with FlowJo software. Transduced cells were gated on the basis of GFP expression. Dead cells were excluded by staining with PI or LIVE/DEAD Fixable Dead Cell Stain Kit (Life Technologies). For RNA-seq, GFP-positive cells were sorted using MoFlo Astrios (Beckman Coulter) or FACSAria II (BD Bioscences).

### Protein extraction and immunoblotting

Cells were washed with ice-cold PBS and resuspended in RIPA buffer to prepare whole-cell lysates. Protein amount was quantified by BCA protein assay reagents (Pierce) and normalized for loading on a 4–12% B-T NuPage gels (Invitrogen). After transfer on nitrocellulose membrane, proteins were stained with antibodies specific for KAP1 and β-actin (Abcam).

### Chromatin Immunoprecipitation

Chromatin was prepared from 10^7^ primary CD4^+^ T cells as previously described (Rowe et al. 2013b). Immunoprecipitations were performed using rabbit polyclonal antibodies specific for KAP1 (Tronolab SY326768, S23470, or Abcam), H3K9me3 (Diagenode), H3K4me1 and H3K27ac (Abcam). ChIP samples were tested in triplicate with SYBR Green mix (Applied Biosystems). Primers (Supplemental Table 2) were designed for an Applied Biosystems 7900HT machine using Primer3Plus (http://primer3plus.com/cgi-bin/dev/primer3plus.cgi). For sequencing, crosslinked cells from three or more donors were pooled and prepared as single ChIP libraries. Total input (TI) and ChIP libraries were prepared as previously described by Santoni de Sio et al. (Santoni de Sio et al. 2012a) using between 2 and 10 ng of material. Sequencing was performed on an Illumina Hi-Seq machine, with each library sequenced in 100-base single-read run. Each ChIP-seq analysis was performed twice in CD4^+^ T cells.

### RNA extraction and quantification

Total RNA was extracted with TRIzol (Invitrogen), purified using miRNeasy kit (Qiagen), treated with RNase-Free DNase (Qiagen) and reverse-transcribed using random hexamers and SuperScript II (Invitrogen). Each cDNA sample was tested in triplicate with SYBR Green mix (Applied Biosystems). Primers (Supplemental Table 2) were designed for an Applied Biosystems 7900HT machine using Primer3Plus or the GetPrime resource (http://updepla1srv1.epfl.ch/getprime) (Gubelmann et al. 2011). Primers specificity was confirmed by dissociation curve analysis and their efficiency was tested by performing reactions with serially diluted samples. Samples were normalized against B2M and RPL13A that were the most stable housekeeping genes in CD4^+^ T cells during activation. RNA-seq in wild-type resting and 72h CD3/CD28-stimulated CD4^+^ T cells was generated with total RNA obtained from two independent donors. Sample libraries were prepared using a TruSeq RNA sample preparation kit (Illumina). RNA-seqs in shLUC, shKAP1-B, and shKAP1-D-transduced cells (CD3/CD28-stimulated, IL-7-treated or rested) were generated with total RNA obtained from three independent donors, after *KAP1* knockdown was verified by RT-qPCR. Sample libraries were prepared using a TruSeq Stranded RNA sample preparation kit (Illumina). Libraries were single-end sequenced on an Illumina Hi-Seq machine with 100-cycles.

### Bioinformatics analyses and statistics

#### ChIP-seq analysis

Reads were mapped to the human genome (hg19) using Bowtie2, with the sensitive local mode (the exact parameters are: bowtie2 -p 6 -t --sensitive-local -x $index -U $reads). KAP1 peaks were called using MACS (Zhang et al. 2008). H3K9me3, H3K4me1 and H3K27ac enriched-islands were identified using SICER (Zang et al. 2009). To detect high confidence binding sites, only peaks with a MACS score above 100 (for KAP1) and with an FDR (as computed by SICER) below 1% (for histone marks) were considered. Correlation analyses between ChIP-seq peaks and genomic features were done with the ChIP-Cor web-based tool (http://ccg.vital-it.ch/chipseq/chip_cor.php). The intersectBed command from the bedtools software (Quinlan and Hall 2010) was used (with the –u parameter) to calculate the intersections between KAP1 and EREs and/or histone marks. Gene annotations were downloaded from Biomart (http://www.ensembl.org/biomart/martview/). The genomeCoverageBed tool from the bedtools software was used to generate coverage files. Those files were converted into bigWig files with the bedGraphToBigWig tool (provided by UCSC) to be visualized on the UCSC Genome Browser. Each displayed track is from a representative biological sample.

#### Public ChIP-seq data

Resting CD4^+^ T cells H3K4me1 and H3K27ac ChIP-seq data were part of the Roadmap Epigenomics Project (http://www.ncbi.nlm.nih.gov/geo/roadmap/epigeneomics/) (Bernstein et al. 2010) and were downloaded from GEO (GSE17312). Activated CD4^+^ T cell ChIP-seq data were generated by Hawkins and colleagues (Hawkins et al. 2013) and were downloaded from the DRAsearch (SRA082670). ChIP-seq tracks for histone marks in mature neutrophils or PBMCs were directly loaded onto UCSC from the Blueprint Epigenome database (http://www.blueprint-epigenome.eu/) or the Roadmap Epigenomics Project, respectively. ChIP-seq track for ZNF274 in GM12878 was generated by Frietze and colleagues (Frietze et al. 2010) and was downloaded from GEO. K562 H3K4me1 and H3K27ac ChIP-seq data were generated by Pradeepa and colleagues (Pradeepa et al. 2016) and were downloaded from GEO (GSE66023). K562 H3K9me3 ChIP-seq data were generated by Salzberg and colleagues (Salzberg et al. 2017) and were downloaded from GEO (GSE71809).

#### RNA-seq analysis

Reads were mapped to the human genome (hg19) using TopHat in sensitive mode (the exact parameters are: tophat -g 2 --no-novel-juncs --no-novel-indels -G $gtf --transcriptome-index $ transcriptome --b2-sensitive -o $localdir $index $reads). The – first-strand option was added for stranded data. Alignment files were then filtered to only contain reads that mapped uniquely to the reference genome. Gene counts were generated using HTSeq-count. For expression analysis, only genes (same for EREs) that had at least as many reads as samples were considered. For repetitive sequences, reads were aligned to a merged repeats track modified from RepeatMasker (see below). EREs counts were generated using the multiBamCov tool from the bedtools software. For both genes and EREs, sequencing depth was corrected using the total number of reads mapped to genes and expression analysis was performed using Voom (Law et al. 2014) as implemented in the limma package of Bioconductor (Gentleman et al. 2004). Genes and EREs were considered as differentially expressed when the fold change between groups was bigger than 2 with a p-value < 0.05, unless specified otherwise. A moderated paired t-test (as implemented in the limma package of R) was used to test significance, correcting p-values for multiple testing using the Benjamini-Hochberg’s method (Benjamini and Hochberg 1995). Files to be visualized on the UCSC Genome Browser were generated as described for ChIP-seq data. Each displayed track is from a representative biological sample.

#### Public RNA-seq data

K562 RNA-seq data were generated by Liu and colleagues (Liu et al. 2017) and were downloaded from GEO (GSE99179).

#### Merged repeats track

All ERE analyses were performed using a merged repeats track generated in-house. The RepeatMasker 4.0.5 (Library 20140131, http://www.repeatmasker.org/species/hg.html) was used as basis. We computed a frequency table of the retroelements surrounding all the different ERV/LTR “-int” elements. To infer the significance of the LTR/int associations we computed the frequencies distributions of the putative LTR for each –int, and performed a Wald test to assign a p-value to each LTR/int pair. The “LTR” was attributed to an “-int” family when the p-value was smaller than 0.001, the frequency higher than 2%, and it was annotated as an LTR. ERV integrants named as “-int” that shared the same name or attributed“LTRs” were merged with their neighboring elements when the distance was shorter than 100 bp. We also kept the fragmented information in our new annotated list. For the final track, the other families of EREs were not modified and were added to the merged ERV/LTR elements.

#### ChIP-seq coverage

Coverage data shown in Supplemental Fig. S2C were extracted from the bigWig files (normalized for sequencing depth) of the different ChIP-seq experiments using the python pybigwig 0.2.7 package. The means of the coverage data for the regions of interest were computed and additionally 95% confidence intervals around the means were calculated.

#### Heatmaps

Tissue histone mark profiles used in Fig. 5 and Supplemental Fig. S11 were downloaded from the NIH Roadmap dataset (Roadmap Epigenomics et al. 2015). The statistical significance of the overlap between KZFP-bound EREs and histone marks was calculated using Bedtools Fisher’s test. Heatmaps were plotted using the heatmap.2 function (R gplots package) and clustering was computed using complete method and euclidean distances.

Tissue gene expression profiles used in Supplemental Fig. S12 were downloaded from the FANTOM5 CAGE dataset (Lizio et al. 2015). Tissues of interest were selected (Supplemental Table 3) and replicates were averaged before plotting. TPM (tags per million) were extracted from the dataset for the genes of interest and the heatmap was plotted using the heatmap.2 function (R gplots package) and clustering was computed using complete method and Pearson distances.

#### Plotting and statistical analysis

Statistical test and plotting were done using R (v3.5.0) or Python programming language (v3.5.1) with the matplotlib python package (v1.5.1) and the seaborn package (v0.7). GraphPad Prism (v8.0) was used for plotting and statistical analysis in Fig. 2D, 3A, 4BCDE, and Supplemental Fig. S2B, S8CD, S9AC and S10BD. Data are presented as mean and standard error of the mean (SEM) of technical or biological replicates (n=3). Statistical significance between datasets was determined by t-test. In case of paired design, we used a paired t-test. Error bars represent SEM. * p-value < 0.05, ** p-value < 0.01, *** p-value < 0.001.

## DATA ACCESS

ChIP-seq and RNA-seq data from this study have been submitted to the NCBI Gene Expression Omnibus (GEO, http://www.ncbi.nlm.nih.gov/geo/) under accession no. GSE81871, GSE81872, and GSE81874.

## ACKNOWLEDGEMENTS

We thank F.R. Santoni de Sio (San Raffaele Telethon Institute for Gene Therapy, Milano) for comments and suggestions, S. Offner for the ChIP-seq libraries preparation, K. Harshman and colleagues (University of Lausanne Genomic Technologies Facility) for high-throughput sequencing, and M. Garcia (EPFL Flow Cytometry Core Facility) for cell sorting. All computing for high-throughput sequencing was done on the Vital-IT cluster. This work was financed through grants from the Swiss National Science Foundation, the European Research Council (ERC 268721; ERC 6946658) and the Personalized Health and Related Technologies to DT. The authors declare no financial competing interest.

## SUPPLEMENTAL FIGURE LEGENDS

**Supplemental Figure S1.**
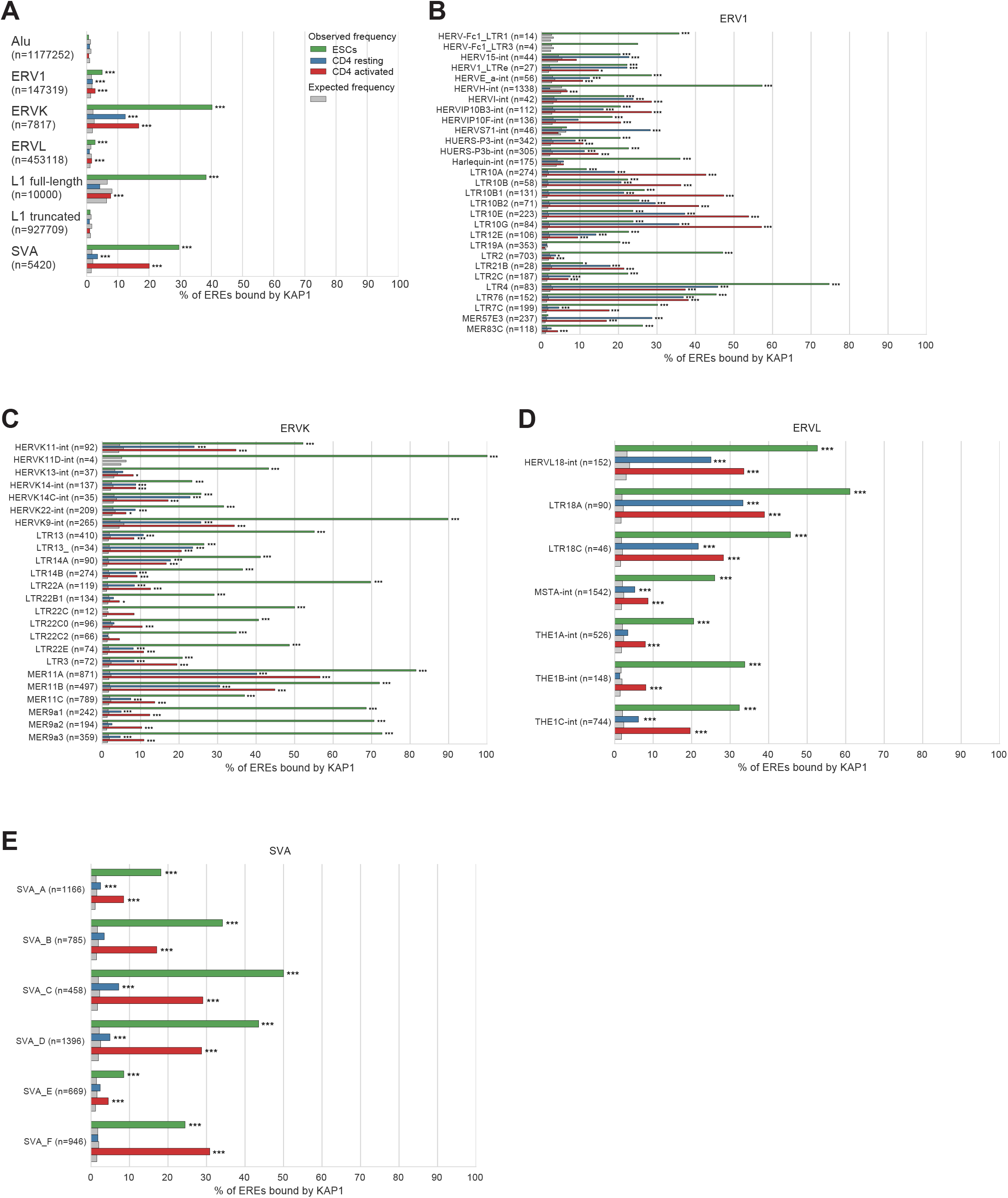
Comparative analysis of KAP1 binding on EREs in ESCs, resting and activated CD4^+^ T cells. (A) Bar plot showing the percentage of ERE integrants on which we detected KAP1 ChIP-seq peaks. For full-length L1 integrants, a length cutoff of 5kb was used. For each family, both the values observed (color bars) or expected randomly (grey bars) are shown. Expected frequencies and statistical significance was calculated using Bedtools Fisher’s test. This experiment represents a biological replicate of the experiment illustrated in Fig. 1B. (B-E) Same as in A, except that they illustrate the percentage of KAP1-bound ERE integrants within subfamilies.

**Supplemental Figure S2.**
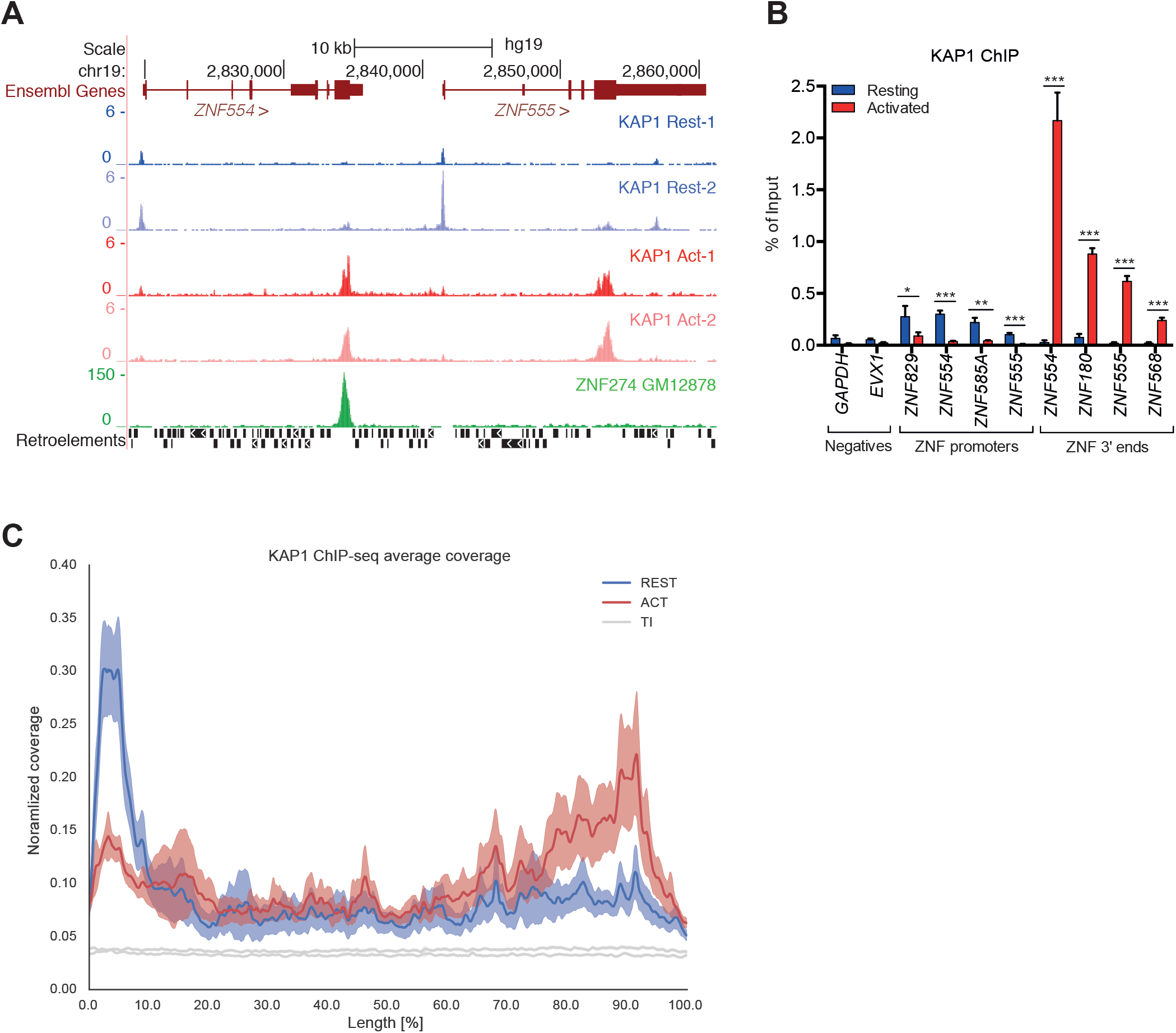
KAP1 binding dynamics at KRAB-ZNF gene loci. (A) UCSC genome browser snapshot of KAP1 ChIP-seq profiles in resting and activated CD4* T cells for a representative ZNF locus on human chromosome 19. Results of duplicate experiments are shown. Displayed tracks include also ZNF274 distribution in GM12878. (B) Eight KAP1 binding sites on promoters or 3’ends of KRAB-ZNF genes, identified by ChIP-seq, were selected to confirm KAP1 enrichment by ChIP-qPCR using independent samples. ChIP values were normalized to their respective total inputs. *GAPDH* or *EVX1* were chosen as negative-control regions. Error bars represent SEM of technical replicates (n=3), *p < 0.05, **p < 0.01, ***p < 0.001, t test. (C) KAP1 ChIP-seq signal normalized for sequencing depth (reads per millions of mapped reads), detected in resting (REST) and activated (ACT) CD4* T cells over the length of KRAB-ZNF genes (lengths were extended of 1kb from the gene start and end coordinates).

**Supplemental Figure S3.**
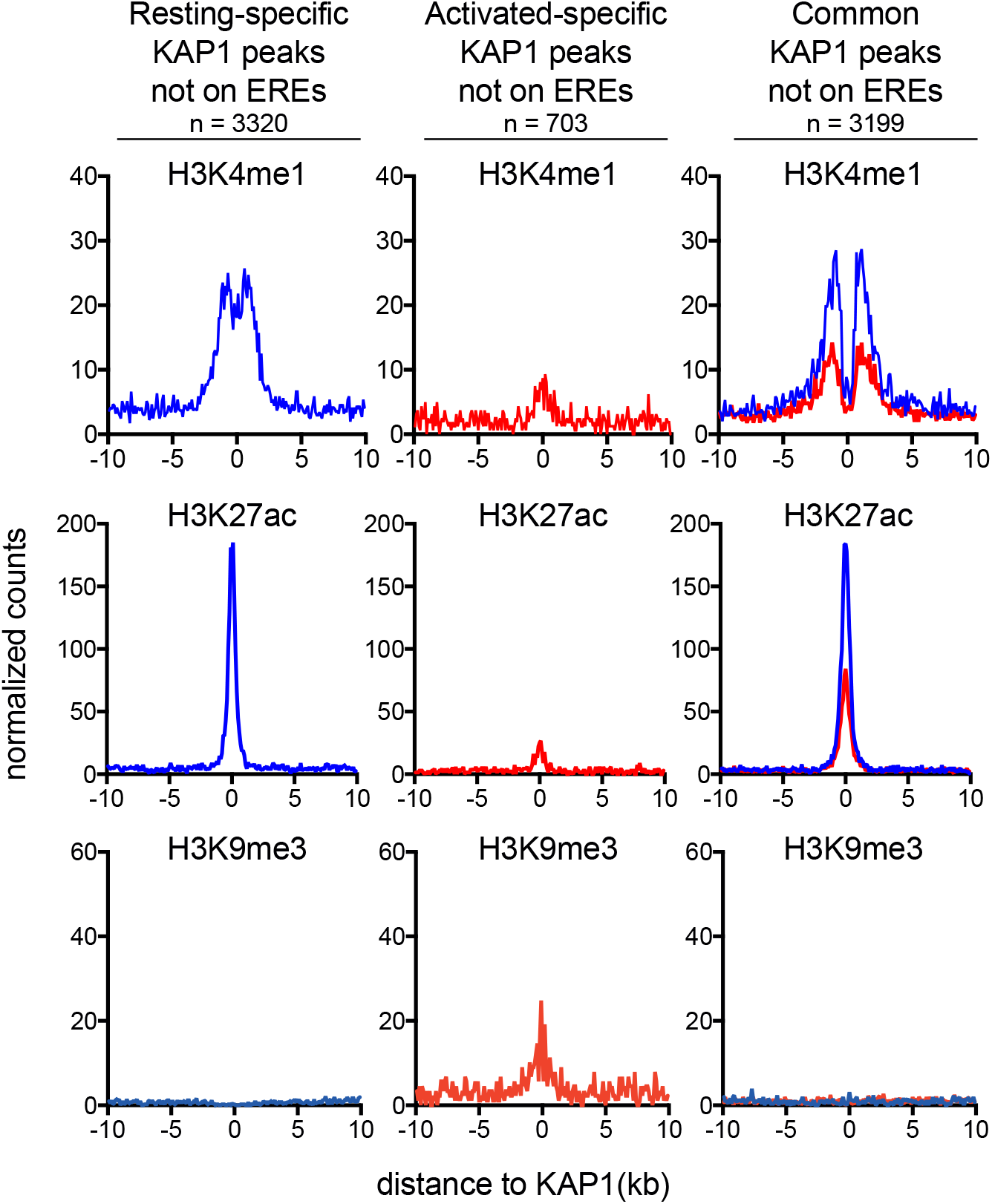
Correlation between KAP1 binding and histone marks at non-ERE loci. Positional relationship between KAP1 peaks not associated with EREs and the indicated histone marks, at 100-bp resolution and over a 10kb window centered on KAP1 ChIP-seq peaks. Left, KAP1 peaks detected in exclusively in resting cells. Middle, KAP1 peaks detected exclusively in activated cells. Right, KAP1 peaks detected in both datasets. The profiles were normalized for the total number of ChIP-seq peaks in each sample.

**Supplemental Figure S4.**
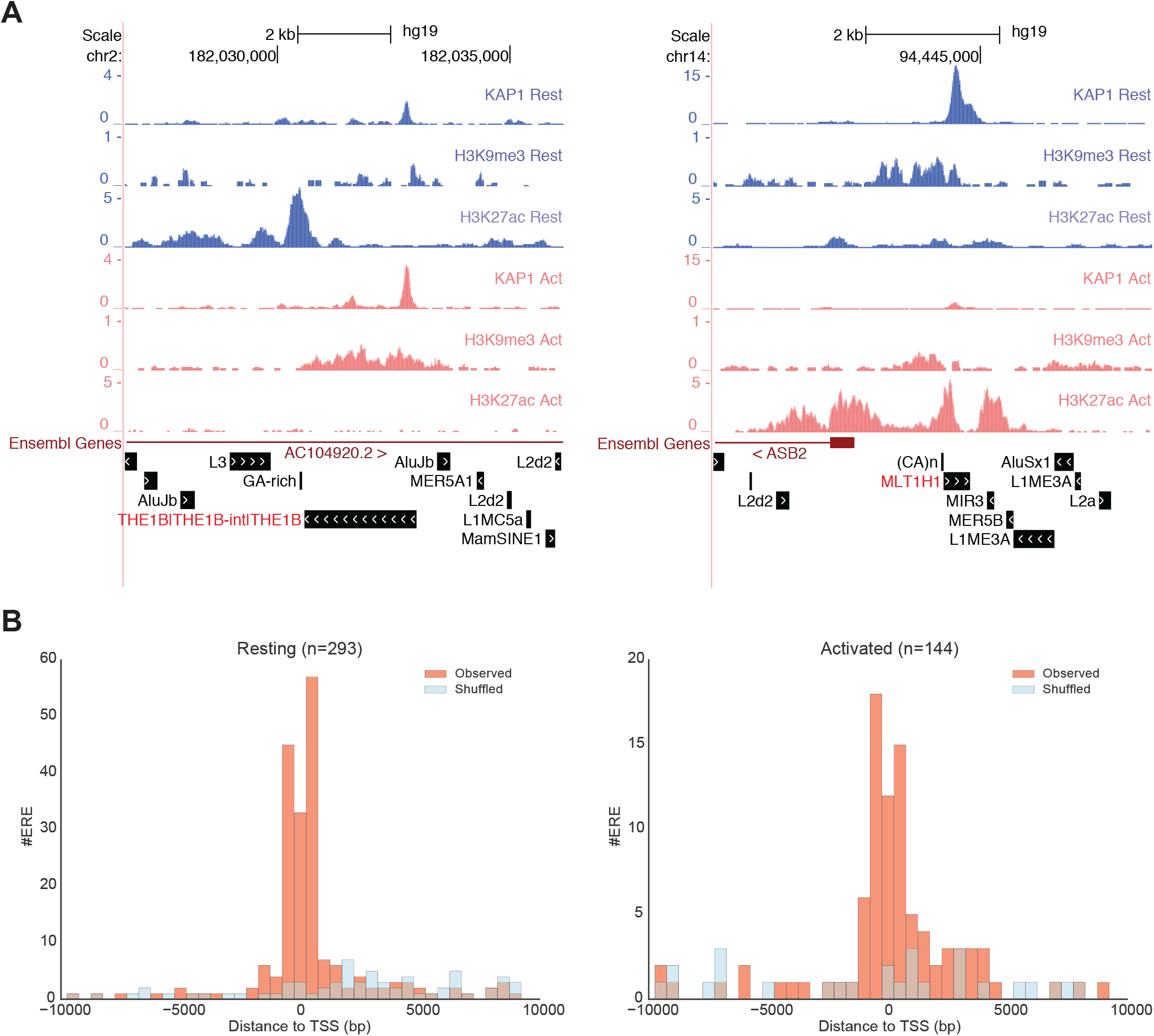
KAP1 binding at ERE-based, activation-dependent regulatory elements. (A) UCSC Genome Browser view of KAP1-bound ERE integrants characterized by a switch from active to repressive (left panel) or from repressive to actve (right panel) chromatin marks following T cell activation. Displayed tracks include KAP1, H3K9me3 and H3K27ac ChIP-seq profiles for resting and activated CD4* T cells. Repeat annotation, downloaded from the UCSC Genome Browser and modified as described in the Methods, is also displayed. EREs integrants on which KAP1 binding is centered are highlighted in red. Results are representative of two independent ChIP-seq experiments. (B) Distance from TSS of resting cell enhancers (left panel) or activated cell enhancers (right panel). For the random set, EREs were shuffled 1000 times within chromosomes. The lists of EREs possessing resting or activated cell enhancer-like signatures were obtained by interrogating them for the presence of KAP1 and H3K9me3-enrichment in one condition and the presence of active H3K27ac mark in the other. EREs shared between the lists were discarded. The total number of identified loci is indicated above each panel.

**Supplemental Figure S5.**
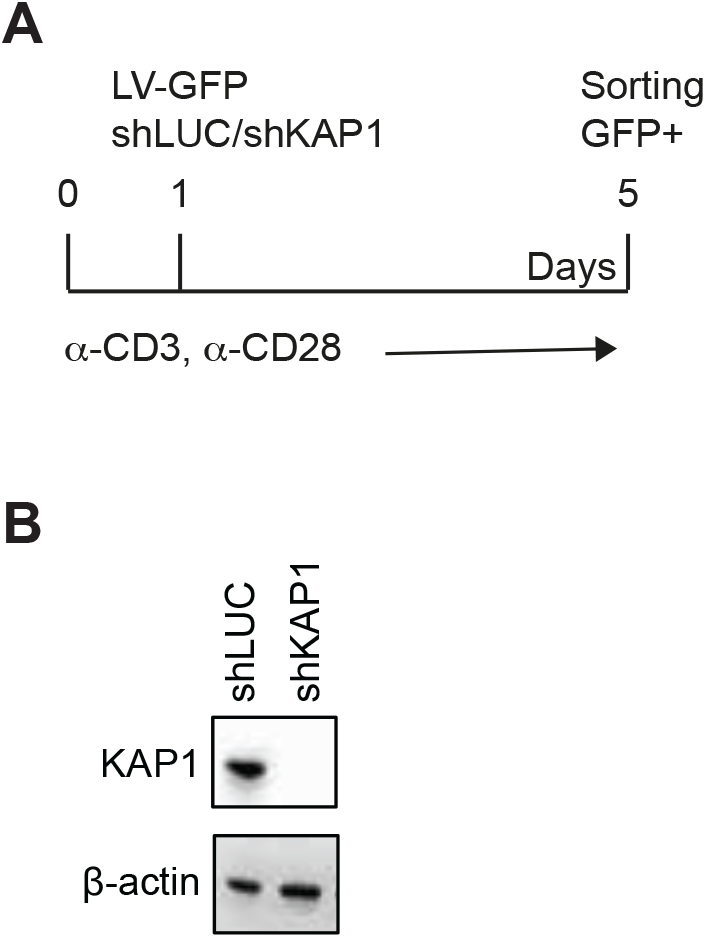
KAP1-depletion in CD4^+^ T cells. (A) Timeline of the experiment. (B) Immunoblot analysis of KAP1 in control (shLUC) and *KAP1* knockdown (shKAP1) CD4* T cells at 5 days after transduction. Beta-actin was used as loading control.

**Supplemental Figure S6.**
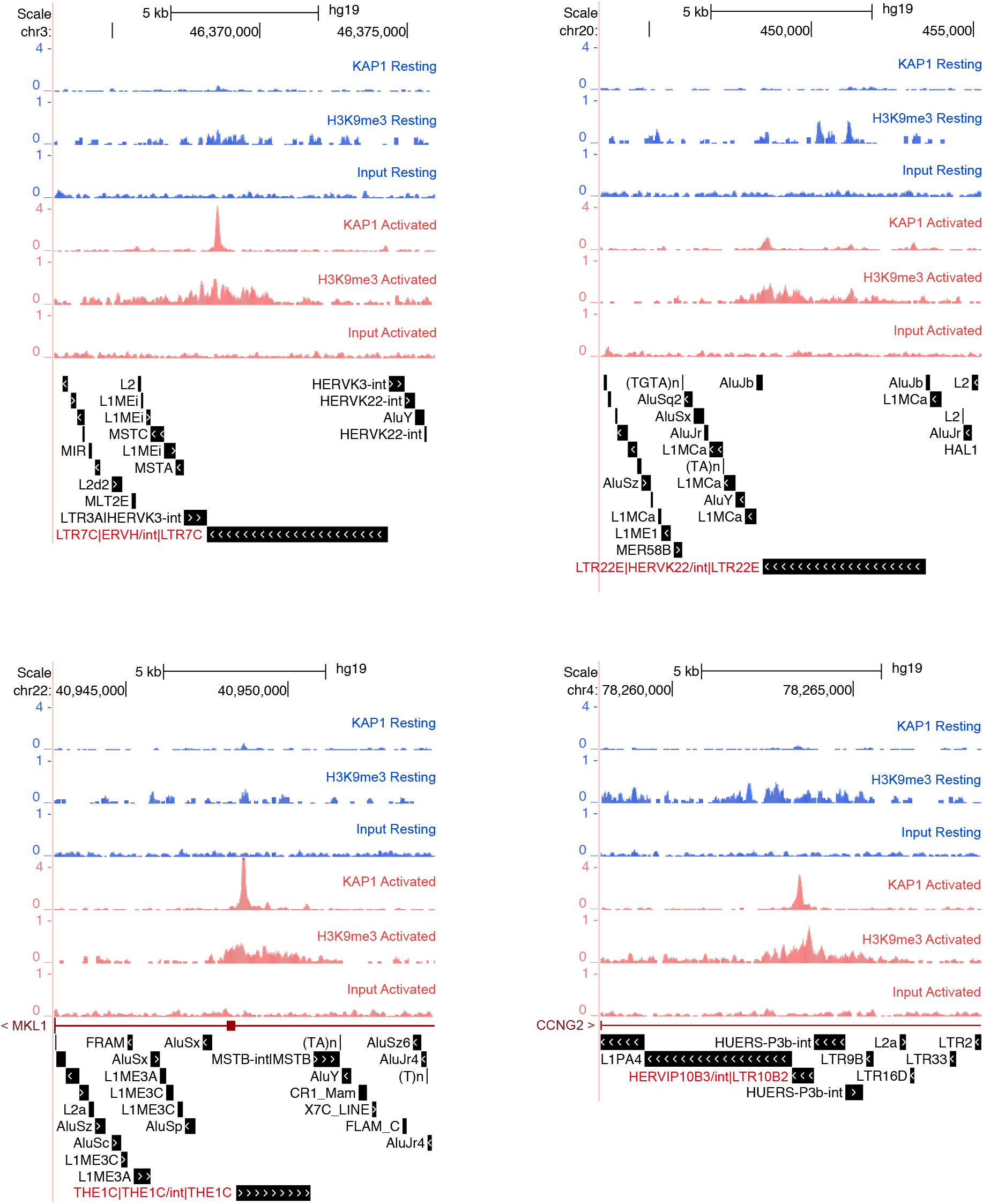
KAP1 recruitment at individual ERE loci. UCSC genome browser snapshots of KAP1 and H3K9me3 ChIP-seq profiles in resting and activated CD4^+^ T cells at individual ERE loci. Results are representative of two independent ChIP-seq experiments. ERE annotation, downloaded from the UCSC Genome Browser and modified as described in the Methods, is also displayed. ERE integrants on which KAP1 binding is centered are highlighted in red.

**Supplemental Figure S7.**
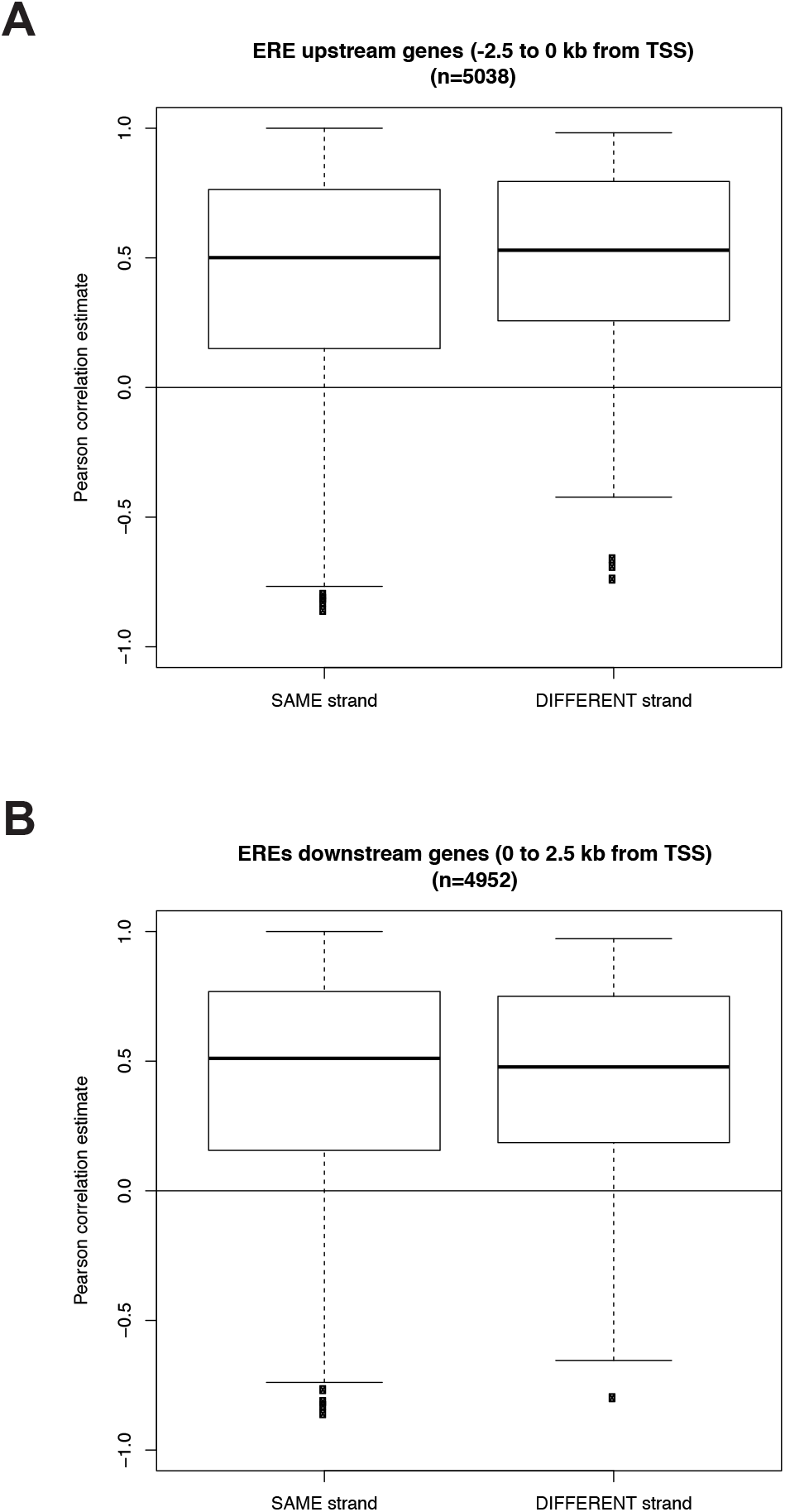
Correlation between expression of EREs and closest genes. Pearson correlation coefficient for EREs located within 2.5kb from the closest genes, encoded in the same or opposite strand. (A) EREs located upstream and (B) EREs located downstream genes.

**Supplemental Figure S8.**
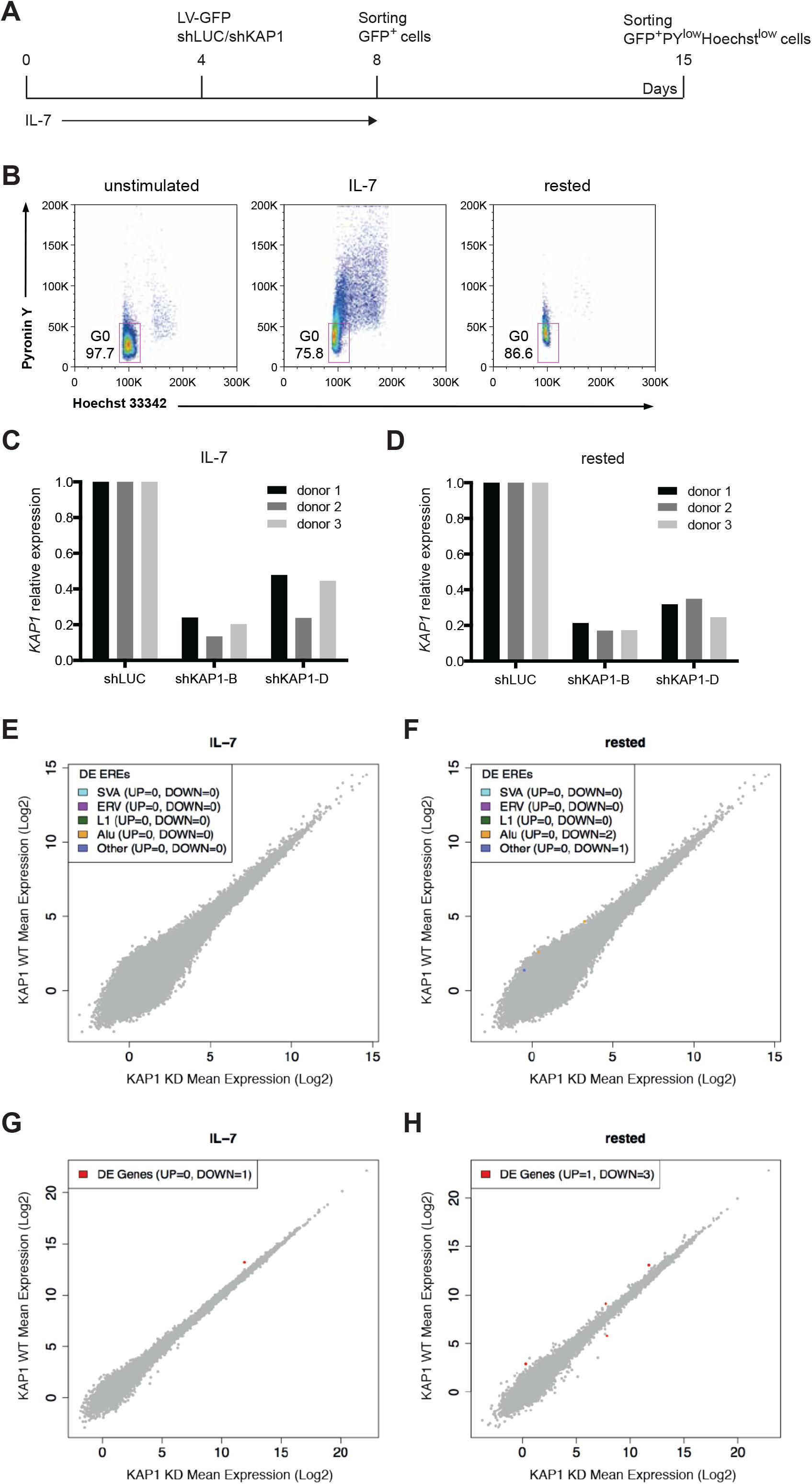
Low transcriptional deregulation in *KAP1* knockdown CD4^+^ T cells without TCR-signaling. (A) Timeline of the experiment. (B) Gating strategy to enrich for quiescent CD4^+^ T cells after IL-7 treatment and rest in culture. Cells were stained for DNA and RNA levels using HOECHST 33342 and Pyronin Y, respectively, and analysed by flow cytometry. Transduced cells were gated on the basis of GFP expression. (C) RT-qPCR validation of KAP1 down-regulation in RNA samples from IL-7 treated or (D) rested CD4* T cells subjected to sequencing analysis. (E) Comparison of ERE expression data from *KAP1* knockdown (shKAP1) and control cells (shLUC) in IL-7 treated or (F) rested CD4* T cells. Results are presented as average normalized counts for each condition. Each ERE integrant is represented by a single data point. Colored data points indicate a significant (P_adj_, < 0.05) and greater than two-fold change in expression. (GH) Same as in E and F but for genes.

**Supplemental Figure S9.**
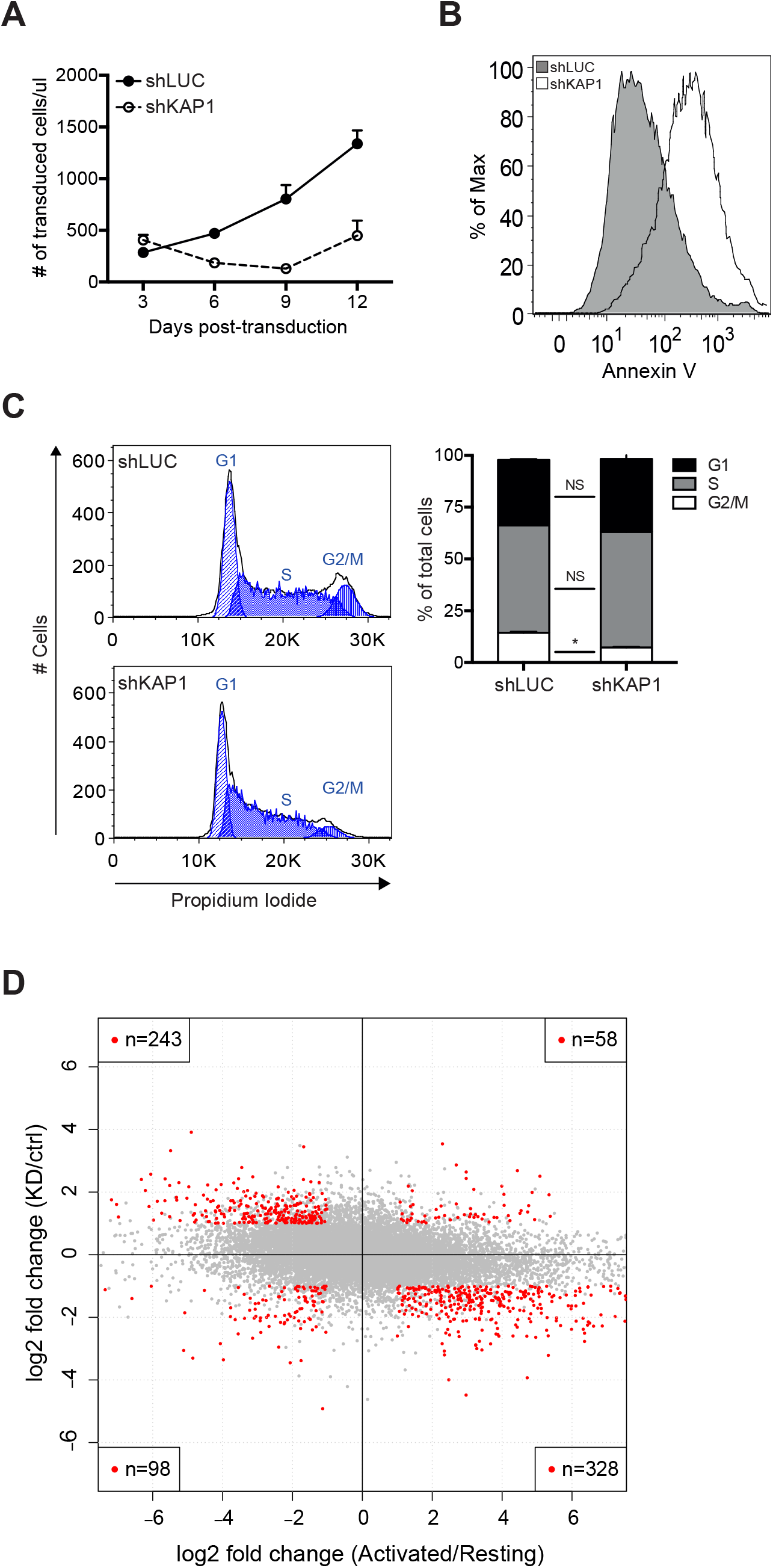
KAP1 depletion counteracts CD4^+^ lymphocytes responses to TCR activation. (A) Number of transduced control (shLUC) and *KAP1* knockdown cells (shKAP1) over time. Cells were enumerated by flow cytometry using CountBright counting beads. (B) Fraction of cells undergoing apoptosis in control and *KAP1* knockdown populations. Apoptosis was measured by Annexin V staining five days after transduction. (c) Cell cycle profile, as assessed by propidium iodide analysis of DNA content, three days after transduction. Left panel: representative cytometry plot. Right panel: cell cycle phase distribution calculated with Watson (pragmatic) model within the Flowjo software package. (D) Comparison of gene expression changes after CD3/CD28-stimulation and *KAP1* down-modulation in CD4* T cells. x axis represents fold change between 72h-activated and resting cells; y axis represents fold change between *KAP1* knockdown and control activated cells at 3 days after transduction with shRNA-expressing LVs. Each gene is represented by a single data point. Red, genes significantly deregulated under both conditions (> 2-fold regulation, p-value < 0.05); gray, genes significantly deregulated only in one or in none dimension. The number of genes significantly deregulated under both conditions is indicated within each quadrant (Pearson correlation estimate = -0.2531; p-value = 2.254e-298).

**Supplemental Figure S10.**
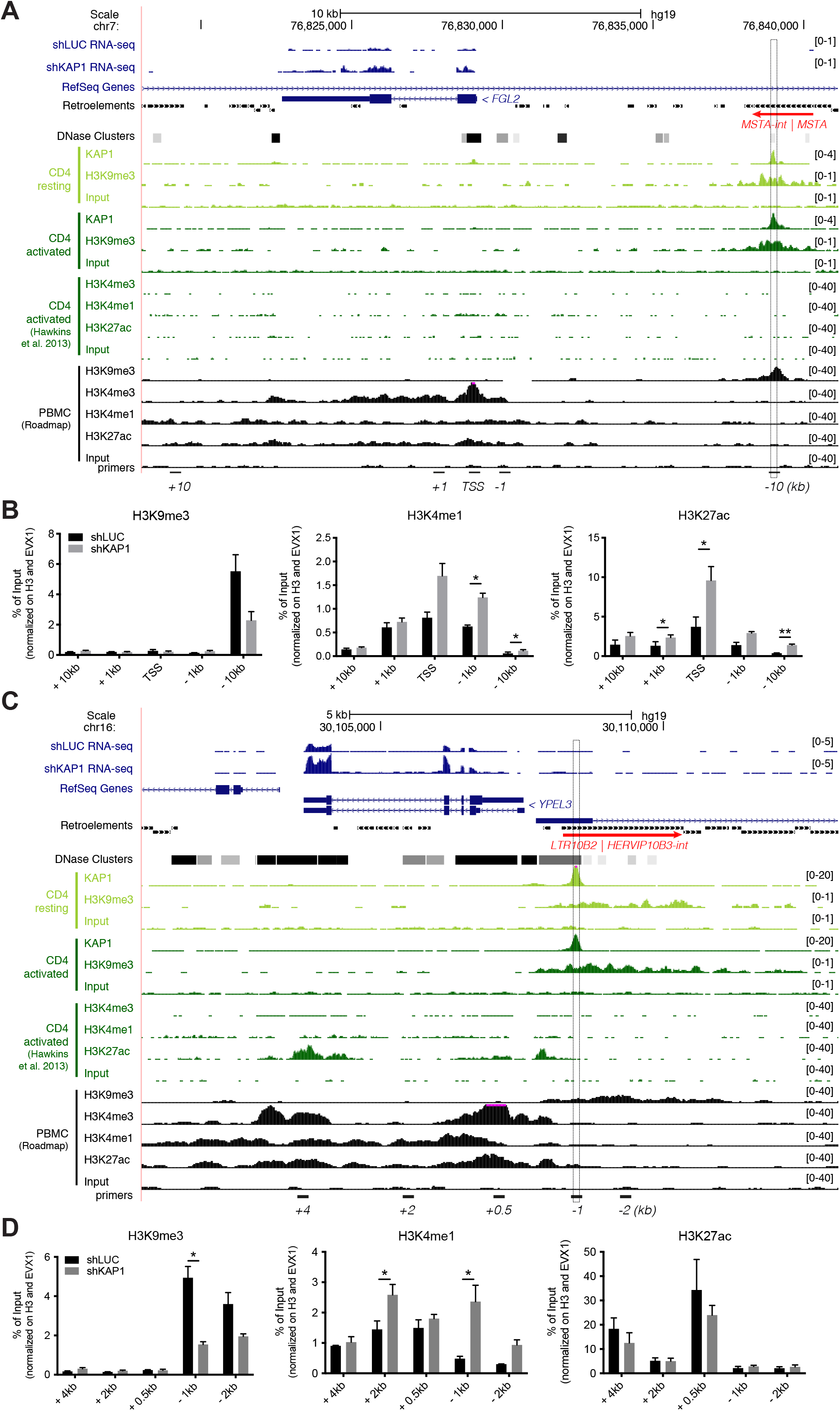
KAP1-regulated expression and histone mark changes at *FGL2* and *YPEL3* loci. (A) UCSC Genome Browser view of *FGL2* locus. Displayed tracks include RNA-seq for control (shLUC) and *KAP1* knockdown (shKAP1) activated CD4* T cells; RefSeq genes and retroelement annotation (modified from UCSC Genome Browser as described in the Methods); DNase cluster (ENCODE); KAP1 and H3K9me3 ChIP-seq tracks for resting activated CD4* T cells; public H3K9me3, H3K4me3, H3K4me1, and H3K27ac ChIP-seq tracks for Th1 activated CD4^+^ T cells and/or peripheral blood mononuclear cells (PBMC); primers used in (B) located at varying distances relative to the TSS. The dashed box highlights the KAP1-bound region. The retroelement on which KAP1 binding is centered is shown in red. (B) ChIP-qPCR analysis of H3K9me3, H3K4me1 and H3K27ac at the *FGL2* locus in control and *KAP1* knockdown cells. Values are normalized to their respective total inputs, to the total H3 protein levels and to *EVX1*. (C) UCSC Genome Browser view of *YPEL3* locus. (D) ChIP-qPCR analysis of H3K9me3, H3K4me1 and H3K27ac at the *YPEL3* locus in control and *KAP1* knockdown cells.

**Supplemental Figure S11.**
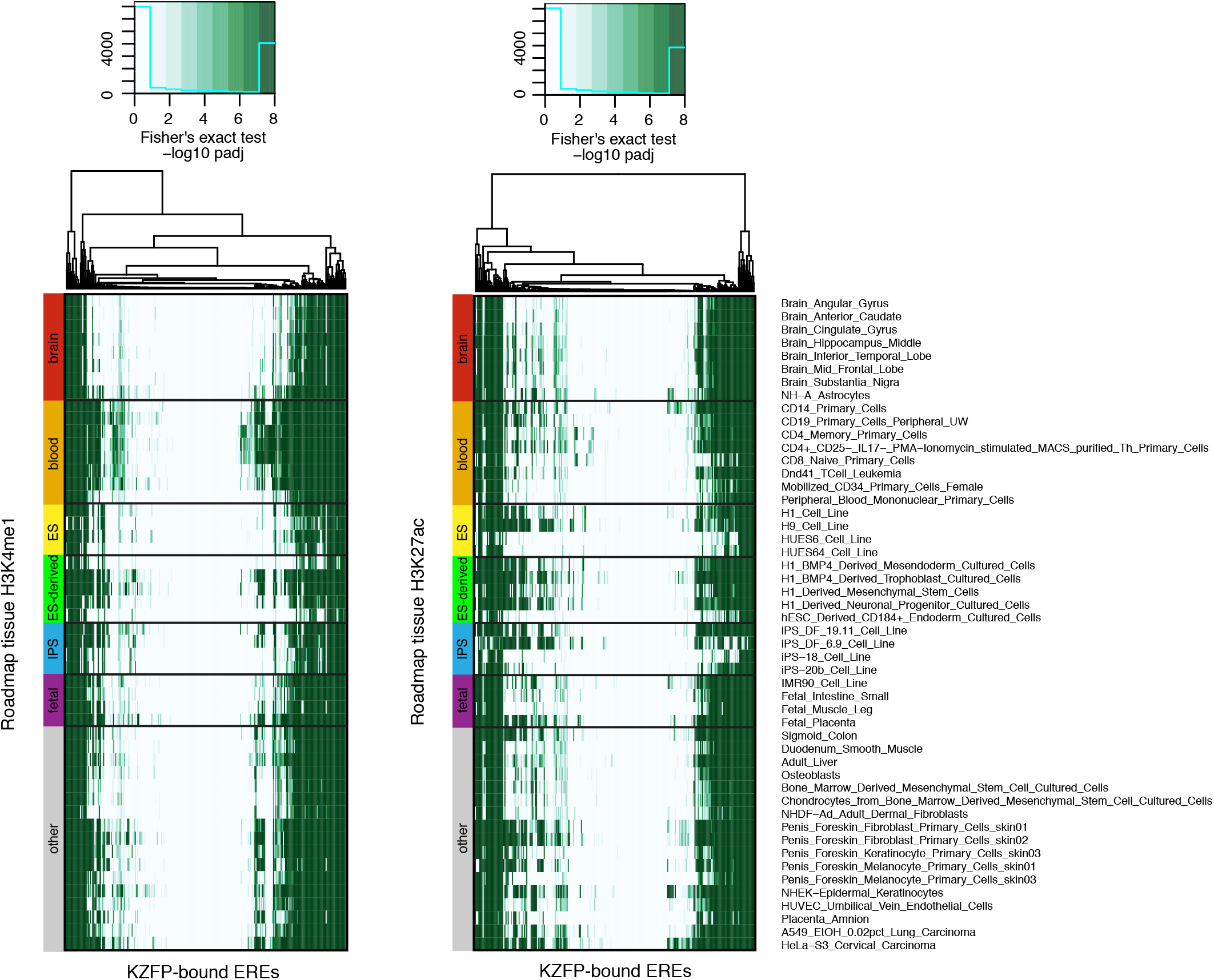
Overlap between KRAB-ZFP-bound EREs and active chromatin marks. (A) Heatmaps showing the Fisher’s Exact test adjusted P-values for the overlap between KRAB-ZFP-bound EREs (X-axis) and H3K4me1 (left panel) or H3K27ac (right panel) from the NIH roadmap dataset (Y-axis). The score is -log10 of Benjamini-Hochberg-corrected P-values.

**Supplemental Figure S12.**
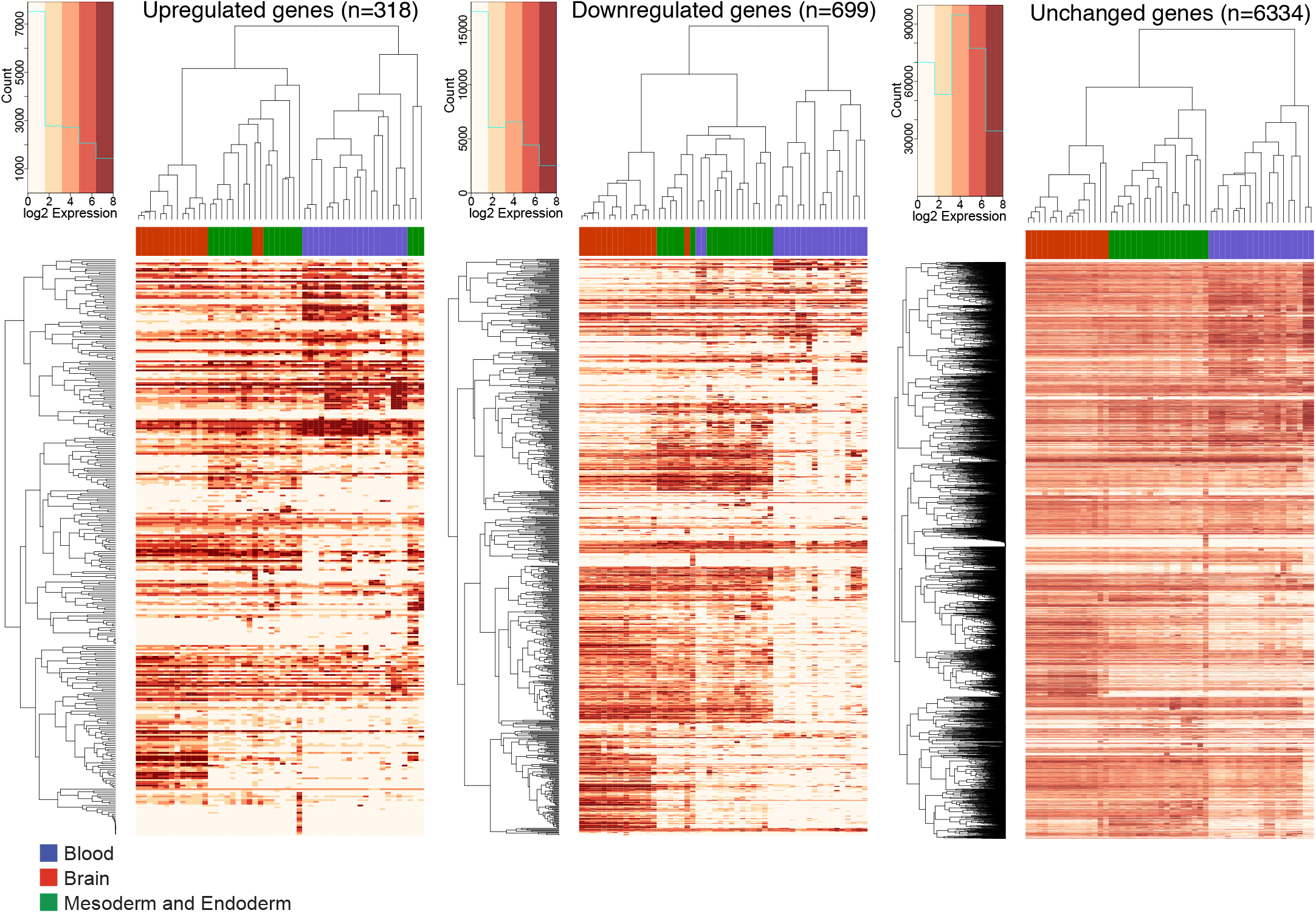
Genes deregulated upon KAP1 depletion display tissue-specific patterns of expression. Clustering of genes upregulated (left panel), downregulated (middle panel), or unchanged (right panel) upon *KAP1* knockdown in activated CD4^+^ T cells on the basis of their expression across a selection of adult tissues grouped into blood (blue), brain (red), and mesoderm and endoderm (green) (for a complete list, see Supplemental Table 3).

## SUPPLEMENTAL TABLES

**Supplemental Table 1.**
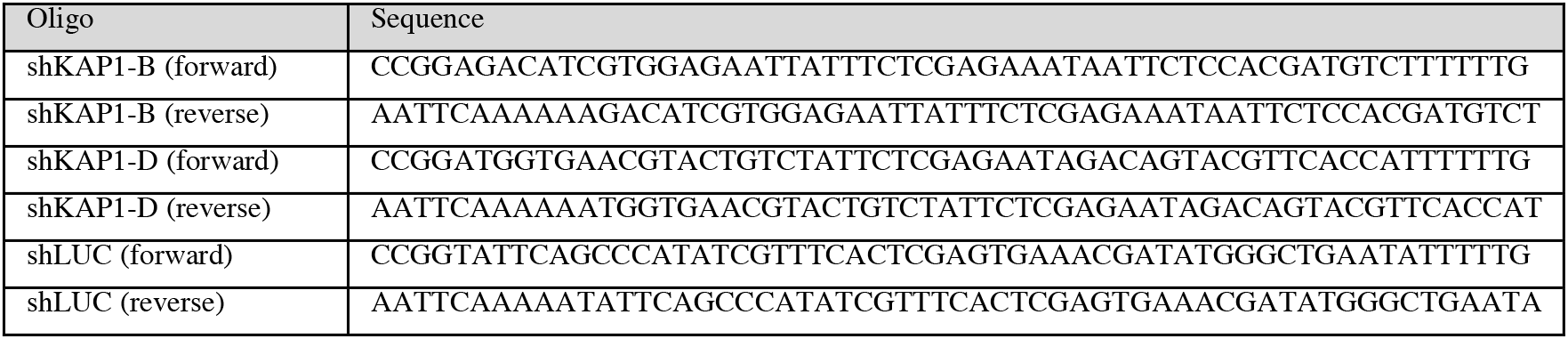
List of shRNA sequences.

**Supplemental Table 2.**
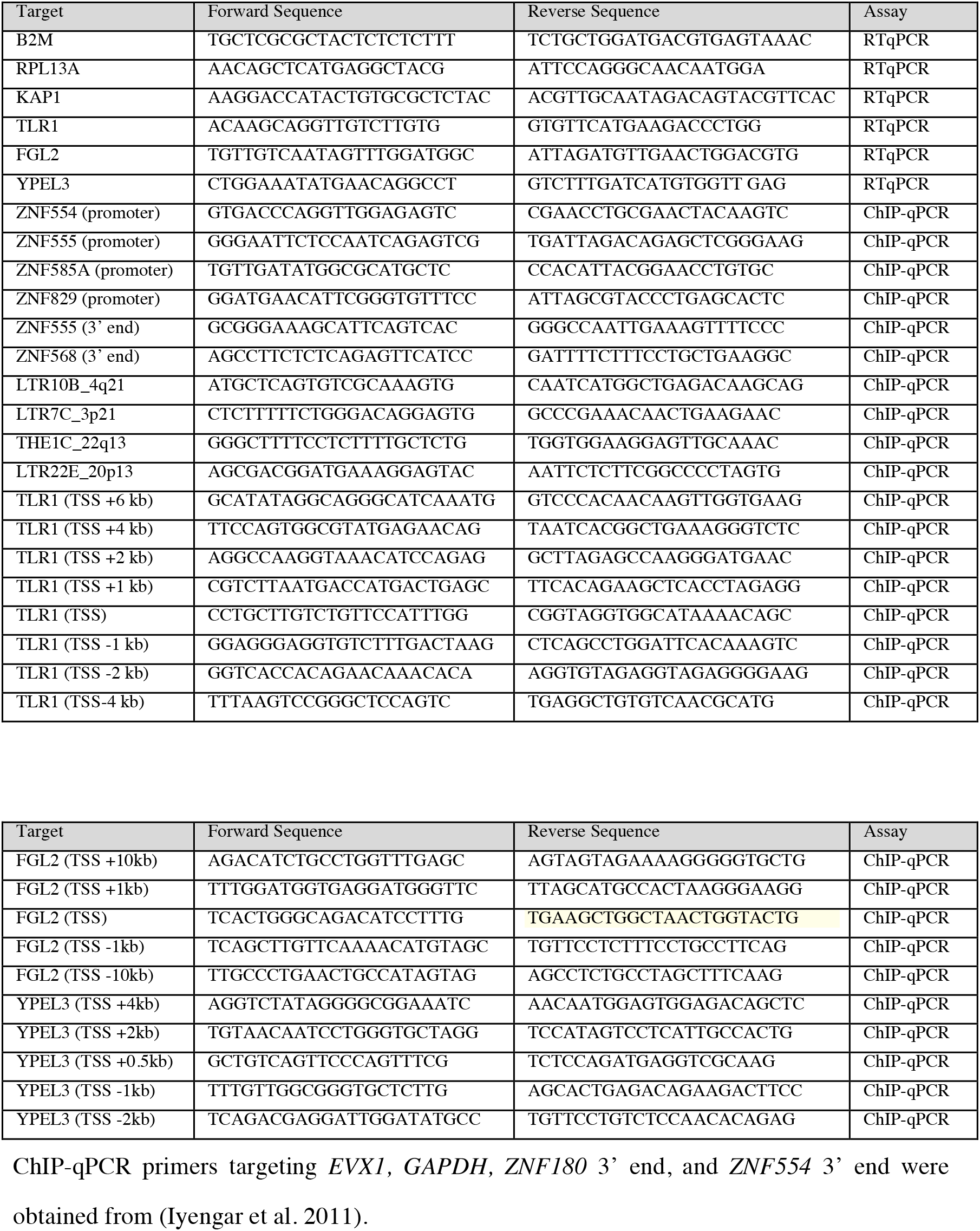
List of primers used for RTqPCR and ChIP-qPCR.

**Supplemental Table 3.** List of transcriptome data generated from the FANTOM5 collection used to analyse tissue specificity of genes.

| Sample | Group |
| --- | --- |
| Basophils | Blood |
| CD14+_Monocytes | Blood |
| CD19+_B_Cells | Blood |
| CD34+_Progenitors | Blood |
| CD4+_T_Cells | Blood |
| CD8+_T_Cells__pluriselect | Blood |
| Dendritic_Cells_monocyte_immature_derived | Blood |
| Dendritic_Cells_plasmacytoid | Blood |
| Eosinophils | Blood |
| Lymph_node_adult | Blood |
| Macrophage_monocyte_derived | Blood |
| Mast_cell | Blood |
| Migratory_langerhans_cells | Blood |
| Natural_Killer_Cells | Blood |
| Neutrophils | Blood |
| Peripheral_Blood_Mononuclear_Cells | Blood |
| Spleen_adult | Blood |
| Thymus_adult | Blood |
| Whole_blood_ribopure | Blood |
| Amygdala_adult | Brain |
| Caudate_nucleus_adult | Brain |
| Cerebellum_adult | Brain |
| Frontal_lobe_adult | Brain |
| Globus_pallidus_adult | Brain |
| Hippocampus_adult | Brain |
| Medulla_oblongata_adult | Brain |
| Occipital_lobe_adult | Brain |
| Parietal_lobe_adult | Brain |
| Pituitary_gland_adult | Brain |
| Pons_adult | Brain |
| Retina_adult | Brain |
| Spinal_cord_adult | Brain |
| Temporal_lobe_adult | Brain |
| Thalamus_adult | Brain |
| Adipose_tissue_adult | Mesoderm and Endoderm |
| Bladder_adult | Mesoderm and Endoderm |
| Colon_adult | Mesoderm and Endoderm |
| Heart_adult | Mesoderm and Endoderm |
| Kidney_adult | Mesoderm and Endoderm |
| Liver_adult | Mesoderm and Endoderm |
| Lung_adult | Mesoderm and Endoderm |
| Ovary_adult | Mesoderm and Endoderm |
| Pancreas_adult | Mesoderm and Endoderm |
| Placenta_adult | Mesoderm and Endoderm |
| Prostate_adult | Mesoderm and Endoderm |
| Skeletal_muscle_adult | Mesoderm and Endoderm |
| Small_intestine_adult | Mesoderm and Endoderm |
| Smooth_muscle_adult | Mesoderm and Endoderm |
| Testis_adult | Mesoderm and Endoderm |
| Thyroid_adult | Mesoderm and Endoderm |
| Trachea_adult | Mesoderm and Endoderm |
| Uterus_adult | Mesoderm and Endoderm |

